# Sequence aligners can guarantee accuracy in almost *O*(*m* log *n*) time: a rigorous average-case analysis of the seed-chain-extend heuristic

**DOI:** 10.1101/2022.10.14.512303

**Authors:** Jim Shaw, Yun William Yu

## Abstract

Seed-chain-extend with k-mer seeds is a powerful heuristic technique for sequence alignment employed by modern sequence aligners. While effective in practice for both runtime and accuracy, theoretical guarantees on the resulting alignment do not exist for seed-chain-extend. In this work, we give the first rigorous bounds for the efficacy of seed-chain-extend with k-mers *in expectation.* Assume we are given a random nucleotide sequence of length ~ *n* that is indexed (or seeded) and a mutated substring of length ~ *m* ≤ *n* with mutation rate *θ* < 0.206. We prove that we can find a *k* = *Θ*(log *n*) for the k-mer size such that the expected runtime of seed-chain-extend under optimal linear gap cost chaining and quadratic time gap extension is *O*(*mn*^*f*(*θ*)^ log *n*) where *f*(*θ*) < 2.43 · *θ* holds as a loose bound. The alignment also turns out to be good; we prove that more than 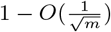 fraction of the homologous bases are *recoverable* under an optimal chain. We also show that our bounds work when k-mers are *sketched*, i.e. only a subset of all k-mers is selected, and that sketching reduces chaining time without increasing alignment time or decreasing accuracy too much, justifying the effectiveness of sketching as a practical speedup in sequence alignment. We verify our results in simulation and on real noisy long-read data and show that our theoretical runtimes can predict real runtimes accurately. We conjecture that our bounds can be improved further, and in particular, *f*(*θ*) can be further reduced.

## 1 Introduction

Since the earliest years of bioinformatics, one primitive task has been sequence *alignment* (Altschul et al., 1990; Smith and Waterman, 1981), which plays a major role in genomic sequencing and phylogenetics. Intuitively, alignment matches together similar parts of two strings, inserting gaps where needed. More formally, alignment is defined by a set of allowed edit operations (e.g. single-character substitutions, insertions, deletions) with associated costs; an alignment is a sequence of those operations transforming one string into another string (or for local alignment, transforming one string into a substring of the other), and the alignment score is the summed cost of those operations (Berger et al., 2021).

Unfortunately, the best guaranteed algorithms for computing alignment (Needleman and Wunsch, 1970; Smith and Waterman, 1981) are quadratic in time-complexity; worse, this bound appears to be tight (Backurs and Indyk, 2018). There exist optimal algorithms with parameterized runtimes based on sequence similarity (Marco-Sola et al., 2021) that are faster in practice, but these still have worst-case quadratic time complexity as a function of the sequence length. Thus, to deal with large volumes of sequencing data (Marçais et al., 2019; Berger et al., 2016), sequence alignment programs employ heuristics (Altschul et al., 1990; Li, 2018; Marçais et al., 2018; Kielbasa et al., 2011; Langmead and Salzberg, 2012; Lipman and Pearson, 1985) without performance guarantees (Medvedev, 2022a) for computational efficiency. All these heuristic algorithms are by necessity fast, achieving empirically sub-quadratic runtimes on real problems. They fail for adversarial examples, but aligners perform well in practice because the sequences being aligned are similar and not pathological examples (Ivanov et al., 2022; Myers, 1986; Ukkonen, 1983; Koerkamp and Ivanov, 2022).

Phylogenetics often makes use of comparative genomics, where aligning together multiple whole genomes allows annotating the ways in which two species have diverged over evolutionary time (Koonin et al., 2000). In genomic sequencing, the alignment task manifests because the sequencing machines are only able to read a small portion of a chromosome at a time, producing short snippets known as *reads* (Alser et al., 2021). It is then incumbent on read-mapping software to determine the likely origin location of that read in the genome, and for already sequenced species, this is usually performed by aligning the read to a known reference genome (Nurk et al., 2022).

Historically, different types of heuristics were used for the two tasks, because aligning a very small substring to a longer string is easier than aligning comparable-size strings. Indeed, the seed-and-extend heuristic, as seen in BWA and Bowtie 2, is preferred for aligning NGS short-reads to genomes instead of aligning genomes to genomes. However, as 3rd-generation long-read sequencing becomes more prominent, the two tasks become more similar and the same heuristics can be used for both (Sahlin et al., 2022)—in this manuscript, we address the *seed-chain-extend* heuristic (distinct from seed-and-extend) used in modern software for both read mapping and whole genome alignment.

### 1.1 Our contribution

Our goal in this manuscript is to close the gap between theory and practice, rigorously justifying the heuristics used in some of the most widely used alignment software. To this end, we turn to the methods of average-case analysis (Szpankowski, 2001), which gives us a way of breaking through the quadratic barrier of alignment (Medvedev, 2022b). Given a random string, we define a substitution model giving rise to a distribution on pairs of inputs and then average our analysis on pairs of strings over this distribution.

Recently, Ganesh and Sy (Ganesh and Sy, 2020) also used probabilistic analysis to show that a heuristic algorithm based on banded alignment can run in *O*(*n* log *n*) time for two length *n* sequences and is optimal with high probability. However, their banded alignment heuristic has not yet seen any usage in practical software, and their analysis is invalid in the case of read mapping as it only pertains to two nearly equal-length strings. Thus we turn instead to the analysis of an empirically battle-tested heuristic for sequence alignment: seed-chain-extend. Seed-chain-extend is a well-established technique for comparative genomics (Abouelhoda and Ohlebusch, 2005; Myers and Miller, 1995), and recently the addition of *sketching* (or subsampling) has made it popular for long-read aligners (Ren and Chaisson, 2021; Chaisson and Tesler, 2012; Sović et al., 2016) including minimap2 (Li, 2018), the primary algorithm our model of seed-chain-extend is based on.

We provide, to the best of our knowledge, the first average-case bounds on runtime and optimality for the *sketched k-mer seed-chain-extend* alignment heuristic under a pairwise mutation model. Our optimality result shows that for large enough k-mer size *k*, the alignment is mostly constrained to be near the correct diagonal of the alignment matrix and that runtime is close to linear when the mutation or error rate is reasonably small. We also show that subsampling 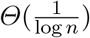 of k-mers asymptotically reduces our bounds on chaining time but not for extension time. Our results give a theoretical justification for both the empirical accuracy and sub-quadratic runtime of seed-chain-extend. A simplified version of our theorem follows.

#### Simplified Theorem 1 (Informal main result; no sketching)

Suppose we are given a uniformly random DNA string of length *n* and a mutated substring of length *m* where each base is substituted with probability *θ.* If *θ* < 0.206 and the longer string is already seeded, then we can choose *k* = *Θ*(log *n*) such that the expected runtime of k-mer seed-chain-extend is *O*(*mn*^*f*(*θ*)^ log(*n*)) = *O*(*mn*^2.43.*θ*^ log(*n*)), and in expectation 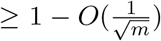 fraction of the homologous bases can be recovered from this alignment.

We will state our models and definitions precisely in Sections 2.1 and 4.4. Our main result, Theorem 1, precisely defines the function *f*(*θ*) < 2.43 · *θ* in the exponent, but 2.43 · *θ* is a convenient upper bound. We can quickly see from this bound that for modest mutation rates, *n*^2.43.*θ*^ is not too large.

## 2 Results

### 2.1 Assumptions and models

#### Mutation model

Let *S* = *x*_1_*x*_2_ … *x*_*n*+*k*-1_ be a random uniform string with *n* + *k* – 1 i.i.d letters on an alphabet of size *σ* for some *k* = *Θ*(log *n*). Let *S*′ = *y_p_*+*y_p_*+_2_ … *y_p_*+*m*+*k*-1 be a substring of *S* of length *m* + *k* – 1 starting at a fixed position *p* + 1 with each character independently mutated to a different letter with probability θ. Although notationally a little confusing at first, the *k* – 1 term ensures *S, S*′ contain exactly *n* and *m* k-mers respectively—k-mers are in many ways the natural unit of measurement, rather than individual characters. We model only point substitutions here and not indels. Independent substitution models have been considered in theoretical work (Blanca et al., 2022; Shaw and Yu, 2022; Ganesh and Sy, 2020), and importantly, also demonstrated to be useful empirically (Chaisson and Tesler, 2012; Ondov et al., 2016; Sarmashghi et al., 2019). Also, while genomes can be repetitive, on the level of k-mers a random model has been shown to be reasonable (Fofanov et al., 2004). We discuss other possible random models in the Discussion section.

#### Modelling seed-chain-extend

A brief overview of seed-chain-extend based alignment is given as follows: first a subset of k-mers in both *S, S*′ are taken as *seeds* and exact seed matches between *S, S*′ called *anchors* are obtained. We only use k-mer seeds in this study, although other types of seeds are possible (Keich et al., 2004; Kiełbasa et al., 2011). An optimal increasing subsequence of possibly overlapping anchors based on some score is then collected into a *chain*, where increasing is defined with the standard precedence relationship (Jain et al., 2022) between k-mer anchors (See Figure 5a and **Chaining** below). The chain is *extended* into a full alignment by aligning between anchor gaps in the chain.

**Model overview:** our model of seed-chain-extend is primarily inspired by minimap2 with a few key differences. It captures the following steps: seeding the *query S*′, matching the k-mers to obtain anchors, sorting the anchors, chaining, and extending. We assume the *reference S* has already been seeded and only the query needs seeding, as in the case of read alignment. For comparing two similar length genomes, the seeding time of either genome is comparable, so the asymptotics will be the same for comparative genomics.
**Runtimes:** non-sketched seeding runtime with a hash table is *O*(*m*), whereas sketched seeding runtime is *O*(*mk*) (discussed in Section 4.5). Letting *N* be a random variable for the number of anchors, matching is *O*(*N* + *m*) by iterating through a hash table, sorting is *O*(*N* log *N*), and (optimal) chaining is *O*(*N* log *N*) (see Section 2.1). Extension is the only step with unknown time complexity. It will turn out that extension and *N* log *N* are the dominating asymptotic terms, so our goal is to bound these terms in expectation. Empirically, it has been shown that chaining and alignment usually take the most time (Kalikar et al., 2022) for read mapping.
**Chaining:** a chain is a sequence of tuples 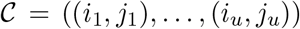 where *i*_ℓ_ and *j*_ℓ_ are the starting positions of the anchoring k-mers on *S* and *S′* respectively, under the convention that *S′* = *y*_*p*+1_ ⋯ so the k-mer labelled (*p* + 1) on *S*′ is actually the first k-mer. The precedence relation *i_ℓ_* > *i*_ℓ-1_ and *j_ℓ_* > *j*_ℓ-1_ must hold for all ℓ, and k-mers can overlap. Our chaining score is the L1 or linear gap cost (Abouelhoda and Ohlebusch, 2005; Li et al., 2020) of the form *u*–*ζ*[(*i_u_*–*i*_1_) + (*j_u_* – *j*_1_)] for *ζ* > 0, which penalizes long chains from distant spurious anchors and is necessary for our proofs when *n* > *m*. The score is sometimes defined equivalently as 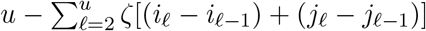. In the language of (Abouelhoda and Ohlebusch, 2005), we let our anchor fragments have length 1, so the k-mers can overlap. While minimap2 v2.22’s default chaining score is different and uses a heuristic banded chaining approach, it does use a linear gap cost (without overlaps) in certain situations, e.g. mapping long contigs (Li, 2021).
**Extension:** we use quadratic time extension between gaps based on any alignment score (e.g. edit distance, affine gap costs (Durbin et al., 1998)) as our optimality criterion only depends on the chain (Section 4.1). We do not extend past the ends of the chain and *do not* use banded alignment (Chao et al., 1992) in this step, unlike minimap2 (Li, 2018).

#### Extension and chaining runtimes

Given sorted anchors, let *T_Chain_* be the time spent finding an optimal chain. *T_Chain_* depends on the objective function (Mäkinen and Sahlin, 2020; Jain et al., 2022; Abouelhoda and Ohlebusch, 2005; Otto et al., 2011). Since our gap costs are linear, *T_Chain_* = *O*(*N* log *N*) where *N* is the number of anchors (Abouelhoda and Ohlebusch, 2005). For extension time *T_Ext_*, let 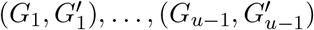 be the size of the gaps for an optimal chain. *G_ℓ_* indicates the length of the substring between the k-mers *i_ℓ_*, *i*_ℓ+1_ on *S* and 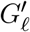 similarly for 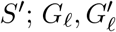 can be zero. The extension time is 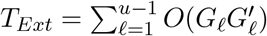. Using the fact that 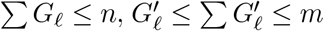, one can get that 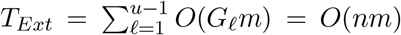. We will show that the expected runtime is better than this upper bound, but it serves as a useful worst case. Since *S, S*′ are random strings, *T_Chain_,T_Ext_,N*, and the alignment itself all become random variables. Our goal will be to bound 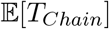 and 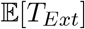.

### 2.2 Theoretical results

First, a few definitions are in order. Recall that |*S*| = *n*+*k* – 1 ~ *n*, and |*S*′| = *m* + *k* – 1 ~ *m* are our string lengths. We define *σ* > 1 to be the size of the alphabet and 0 < *θ* < 1 to be the probability a base mutates. Our theory holds for any alphabet size, but we use *σ* = 4 for specifying numerical constants. Let log = log_*σ*_ with base *σ* and ln be the natural logarithm. Let *k* = *C* log *n* for a fixed *C* > 0 so that key quantities can be expressed interchangeably as *σ^k^* = *n^c^* and (1 – *θ*)^*k*^ = *n^-Cα^* where *α* = – log_*σ*_(1 – *θ*) > 0. We define the actual goodness of the chain in terms of *recoverability* in Section 4.1; it measures the fraction of homologous bases under our mutation model that could potentially be recovered by extending through an optimal chain and *only depends on the chain*, not the actual alignment.

#### Theorem 1 (Main result; no sketching)

*Under our model of seed-chain-extend in Section 2.1, assume θ* < 0.206, *let α* = – log_4_(1 – *θ*), *and pick any* 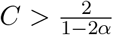. I*f m* = *Ω*(*n*^2*Cα+ε*^) < *n for some arbitrary ϵ* > 0, *letting k* = *C log n and* 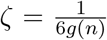 *where* 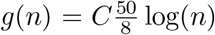 ln(*n*)*n^Cα^, the expected running time is O*(*mn^Cα^* log(*n*)) *for extension and O*(*mn*^-*Cα*^ log *m*) *for (optimal) chaining. Cα can always be chosen to be* 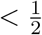 *and the expected recoverability of any optimal chain is* ? 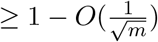.

Equivalently, 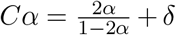 for any 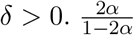 is plotted in Supplementary Figure 7; it is convex and thus leads to the bound *Cα* < 2.43·*θ* stated in Section 1.1. E.g., if |*S*| = |*S*′| and *θ* = 0.05, which is approximately the level of divergence seen between human and chimpanzee genomes (Britten, 2002), 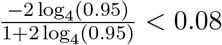, so for *Cα* = 0.08 the running time is *O*(*n*^1.08^ log(*n*)). Even for relatively large values of *n*, say the entire human genome which has size of approximately 3 billion, *n*^1.08^ is essentially linear as (3, 000, 000, 000)^0.08^ ~ 5.73.

Our proof follows in three main steps. First, we bound the first and second moments of the random variable *N*, which denotes the number of anchors, implying that chaining is fast. Second, we use concentration inequalities for sums of *dependent* random variables and exploit the structure of chaining to show that with high probability, an optimal chain does not deviate much from the chain with only “homologous anchors” (Figure 5a). The failure probabilities will be on the order of 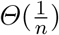, allowing us to also bound everything in expectation. Lastly, we bound the expected runtime of extension through gaps between homologous anchors.

Asymptotically, the chaining runtime is smaller than extension runtime. However, the implied constants can be much smaller for the extension term (Kalikar et al., 2022), so it is practically useful to reduce the runtime of chaining via sketching. Let 0 < 1/*c* ≤ 1 be the expected fraction of selected k-mers for a sketching method. We use the *open syncmer* k-mer seeding method (Edgar, 2021) which has a useful mathematical property (Theorem 7), giving:

#### Theorem 2 (Sketched result)

*In addition to the hypotheses outlined in Theorem 1, let c* = *O*(log *n*) < *k and* 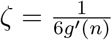 *instead where* 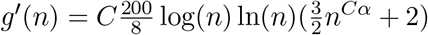. *For open syncmer sketched seed-chain-extend, the expected running time is* 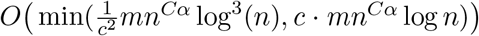 *for extension and* 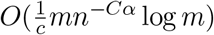 *for chaining. The expected recoverability of any optimal chain is* 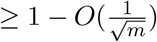.

This shows that the asymptotic upper bound on extension runtime is the same even if we let *c* grow with *n* like *c* = *Θ*(log *n*) < *k*, leading to the following conclusion: *sketching can reduce chaining time without increasing extension time much*. Other seeding schemes used, e.g. minimizers (Sirén et al., 2021; Rautiainen and Marschall, 2020; Li, 2018; Colquhoun et al., 2021) or FracMinHash (Irber et al., 2022) behave differently, but our techniques provide intuition and in some cases can be extended (See Section 4.5).

### 2.3 Simulated genome alignment experiments

Although the model of seed-chain-extend we consider in the theory is based off of real aligners (e.g. minimap2), real aligners implement many additional tricks to make things work better. As such, to empirically validate our theory, we implemented a basic version of sketched seed-chain-extend and tested it on simulated random sequences with independent point substitutions. For the chaining step, we implemented an AVL tree based max-range-query method (Mäkinen et al., 2015; Li et al., 2020). For the extension step, we used a standard dynamic programming (DP) algorithm implemented in rust-bio (Köster, 2016).

We let *k* = *C* log *n* where 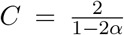, and do seed-chain-extend on two length *n* sequences in this section, but we do an additional experiment for substrings of length *m* = *n*^0.7^ + 100 in *k* Supplementary Figure 10. As *k* cannot be fractional, we let 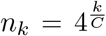 for varying integer values of *k*. We will set *c* = *k* – 7 = *C* log(*n*) – 7 which *grows with n*. The constant 7 is arbitrary but chosen sufficiently large that so s-mers in the open syncmer method are not so small as to make the method degenerate. We found that recoverability was always quite high and that breaks did not occur very often in actual simulations; we show this in Supplementary Figure 9. Thus the primary focus of the empirical results will be runtime. We used 50,000 iterations for every data point.

#### Accuracy of asymptotic extension runtime predictions

We first empirically investigate our upper bound on expected extension runtime, which is *O*(*n*^1+*Cα*^ log(*n*)) for both sketched and non-sketched extension when *c* = *Θ*(log*n*). Assuming that the runtime is simply 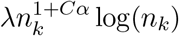 for some fixed constant λ, we can predict the ratio of the runtimes as 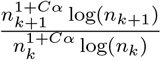. Of course, this is incorrect for small *n*, as smaller terms may dominate runtime, but we expect it to be reasonably accurate for large *n*. We plot the empirical and predicted ratios of extension runtimes in Figure 1 (a).

**Fig. 1:**
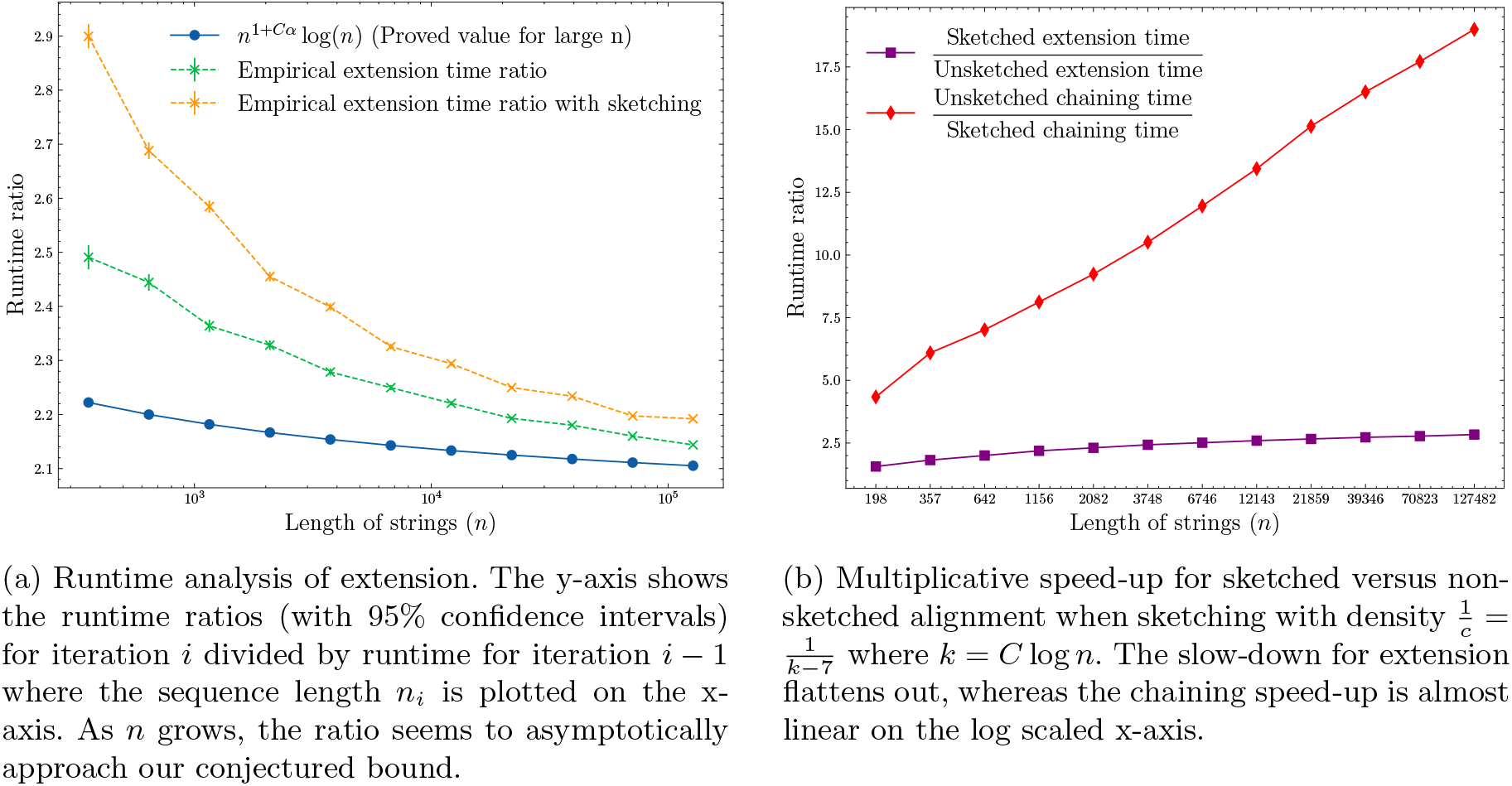
Runtime results on seed-chain-extend between for two sequences of length *n* and 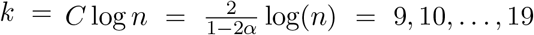 where *θ* = 0.10 and *α* = – log_4_(1 – *θ*). In (a), both sketched/non-sketched extension ratios approach the predicted 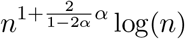 runtime ratio. In (b), chaining speed up dominates extension slow down as predicted by our theory.

Figure 1 (a) shows that our upper bound looks reasonable for both sketched and non-sketched cases for *θ* = 0.10. In Supplementary Figure 8, we show the same plots for *θ* = 0.05. The empirical results never cross the predicted ratio, but this is not too surprising as the predicted ratio is only approximate. Importantly, the empirical extension runtime ratios slope downwards, which is in agreement with the prediction.

#### Sketching with 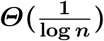 density gives favourable runtime tradeoffs

One of our key results in Theorem 2 is that the upper bound on asymptotic runtime of extension does not depend on the density as long as 1/*c* < *k*, but chaining speed-up scales multiplicatively with *c*. In Figure 1 (b), we plot the multiplicative speed-up and slow-down of chaining and extension runtimes. This figure shows why sketching is so effective in practice with respect to runtimes; the maximal extension slow-down plateaus to ≤ 3 times in this case, whereas the chaining speed up is linear as the string grows exponentially. Therefore it is worth sketching aggressively to reduce runtime if chaining is slow. One should however still be careful of the sensitivity loss due to sketching in practice (Shaw and Yu, 2022).

In practice, because extension is heavily optimized (Marco-Sola et al., 2021; Farrar, 2007; Suzuki and Kasahara, 2018) and repetitive k-mers lead to more anchors, chaining can be a bottleneck even though it is asymptotically faster in runtime. In fact, we tried using WFA (Marco-Sola et al., 2021), a highly optimized algorithm for extension instead of the generic DP algorithm and found it was ~ 60 times faster than chaining without sketching for *n* ~ 2, 800, 000, *k* = 23, *θ* = 0.05 in our implementation.

### 2.4 Real nanopore read alignment experiments

To test the applicability and generalizability of our theorems to real, non-simulated data, we performed an experiment by aligning real long-read data from Oxford Nanopore Technologies (ONT) using seed-chain-extend. We use the sketched seed-chain-extend aligner in the simulated experiments with a slight modification (see the next section). Notably, these reads are smaller than the genome length. Importantly, our two main assumptions with respect to uniform random strings and point substitutions are violated in this setup: real genomes are not uniformly random strings, and ONT reads contain a significant amount of *indel errors* (Delahaye and Nicolas, 2021).

Our data set consists of five reference genomes and corresponding long-read data sets. The references consist of a viral genome (SARS-CoV-2), a bacterial genome (E. coli), a fungi genome (M. oryzae), an insect genome (D. melanogaster), and a human genome (H. sapiens). For each genome, we aligned a corresponding set of publicly available ONT long-reads which may have different length and error distributions. The data sets can be found in Supplementary Table 1.

We are interested in testing the predicted results as a function of *m* (the read length) and *n* (the genome size). We let 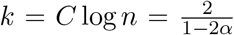 as before. Because *α* and *C* in our results depend on the error rate *θ*, we first aligned each read set to the genomes with minimap2 and retained only reads with gap-compressed identity > 94.5% and < 95.5% so that *θ* ≈ 0.05 is fixed. In addition, we only used reads for which minimap2 aligned > 90% of the read, as to avoid benchmarking on unalignable reads.

#### Practical linear-gap cost consideration

We use almost the same seed-chain-extend algorithm as in Section 2.3 with fixed density 1/7 sketching (i.e. not scaling with *n*), except we apply one change: we found that the predicted linear-gap cost *ζ*(*n*) given by our theorems, while asymptotically appropriate in theory for large *n*, was too small for reasonable *n*. Therefore, we multiplied *ζ* by 1000.

This discrepancy is because the bounds we use in our proofs do not have optimized constants. The exact (non-asymptotic) value of *ζ* is only used in proving the recoverability result, and our change does not modify *ζ*’s asymptotic behavior, which is used in other parts of our proofs. Thus our runtime results are still technically valid, while our experiments below will also suggest that the recoverability result is also still valid.

#### Evaluating model assumptions

We first introduced the following filters to our basic seed-chain-extend alignment algorithm for computational reasons:

1. We filtered reads where the number of anchors is > 10 times the number of bases. This occurs due to extremely repetitive k-mers, and can cause the chaining step to stall.
2. We did not consider reads where the chains had gaps of length > 10000 bp, which could occur due to structural variations or failure of chaining and can cause extension to stall.

The two filters also measure how badly our model assumptions, with respect to independent substitutions and uniform random strings, are violated. We report the number of reads that fail filters (1) and (2) in Table 1. Only a small fraction of reads fail the 10000 bp gap filter, but the repetitive k-mer filter is violated more frequently as the genomes become more complex. Still, > 80% of the human reads and > 97% of the D. melanogaster reads are not deemed too repetitive. Mapping to repetitive regions is an active area of research (Jain et al., 2020) which is out of the scope of this article, but effective heuristics such as repetitive k-mer masking (Li, 2018; Sahlin, 2022) exist.

**Table 1:** Counting the number of reads which did not pass our gap filter and repetitive k-mer filter. These two statistics measure how badly our model breaks down with respect to repetitiveness, large variation, and chaining failure. The total number of reads only counts a portion of the original reads that pass our filters, i.e. 90% of the read is aligned by minimap2 and the sequence divergence is 5% ± 0.5.

|  | # reads with<br>>10 kb gap in chain | # reads with<br>number of anchors ><br>10 times number of bases | Total # of reads |
| --- | --- | --- | --- |
| SARS-CoV-2 | 0 (0%) | 0 (0%) | 1692 |
| *E. coli* | 0 (0%) | 0 (0%) | 8223 |
| *M. oryzae* | 21 (0.39%) | 3 (0.06%) | 5424 |
| *D. melanogaster* | 25 (0.56%) | 116 (2.61%) | 4439 |
| *H. sapiens* | 11 (0.07%) | 3004 (19.48%) | 15419 |

#### Extension and chaining runtimes are well predicted by theory

In Figure 2, we plot the times of chaining and extension and perform a robust linear regression using the Siegel estimator (Siegel, 1982) as a function of the read length *m*. For this plot specifically, we only timed alignments where the read was > 90% aligned by our seed-chain-extend implementation. For each plot, the genome size *n* is held fixed. For fixed *n*, all runtimes are well-approximated by a linear function in *m* as predicted by our theory, although chaining times for the human genome have higher variance due to the presence of repetitive k-mers.

**Fig. 2:**
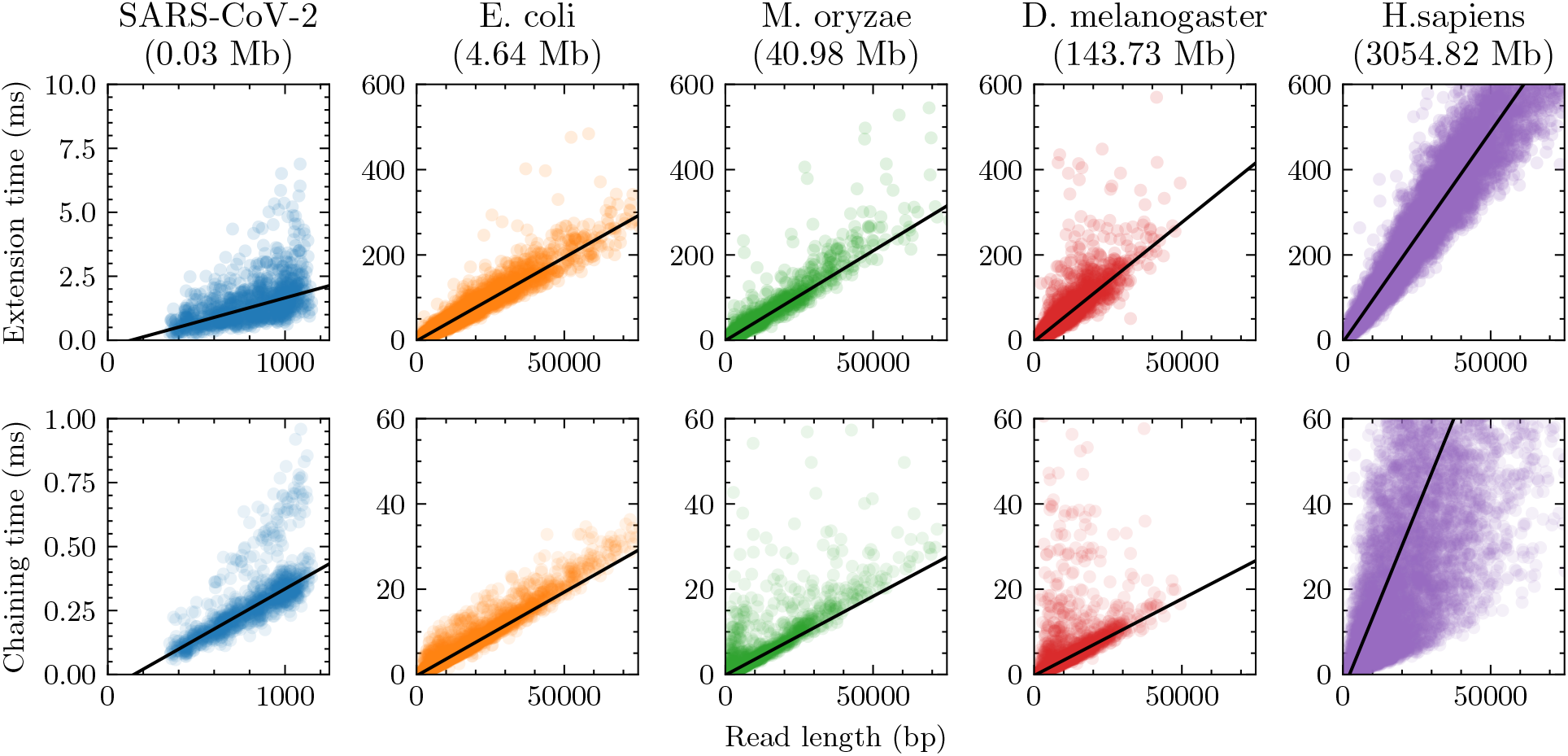
Extension and chaining time for real ONT reads with approximately 5% sequence divergence. k-mer size was chosen to be *k* = *C* log *n* for reference length *n* and 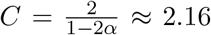 with error parameter *θ* = 0.05. Exact data sets are described in Supplementary Table 1. Note the scaled axes for the SARS-CoV-2 reads, which were much smaller than the other data sets. The well-fit linear regression lines indicate essentially linear runtime in read length with fixed reference length *n* (i.e. fixed *k* = *C* log *n* and constant *C*), although larger human chaining time variance is due to repetitive k-mers.

In Figure 3, we plot the slopes of the regressions for extension time obtained from the first row of Figure 2, which measures the dependence on *n* given fixed *m*. This dependence is reasonably fit by a log*n* function (R^2^ = 0.766), so *O*(*m* log *n*) is not a bad approximation for the runtime in practice. However, our theory leads to a *n^Cα^* log *n* dependence, which comes from the big O extension runtime. *Cα* ~ 0.08 when *θ* = 0.05 in our setup, so by using a *n*^0.08^ log *n* function instead, we end up getting a better fit (*R*^2^ = 0.928), as predicted by our theory.

**Fig. 3:**
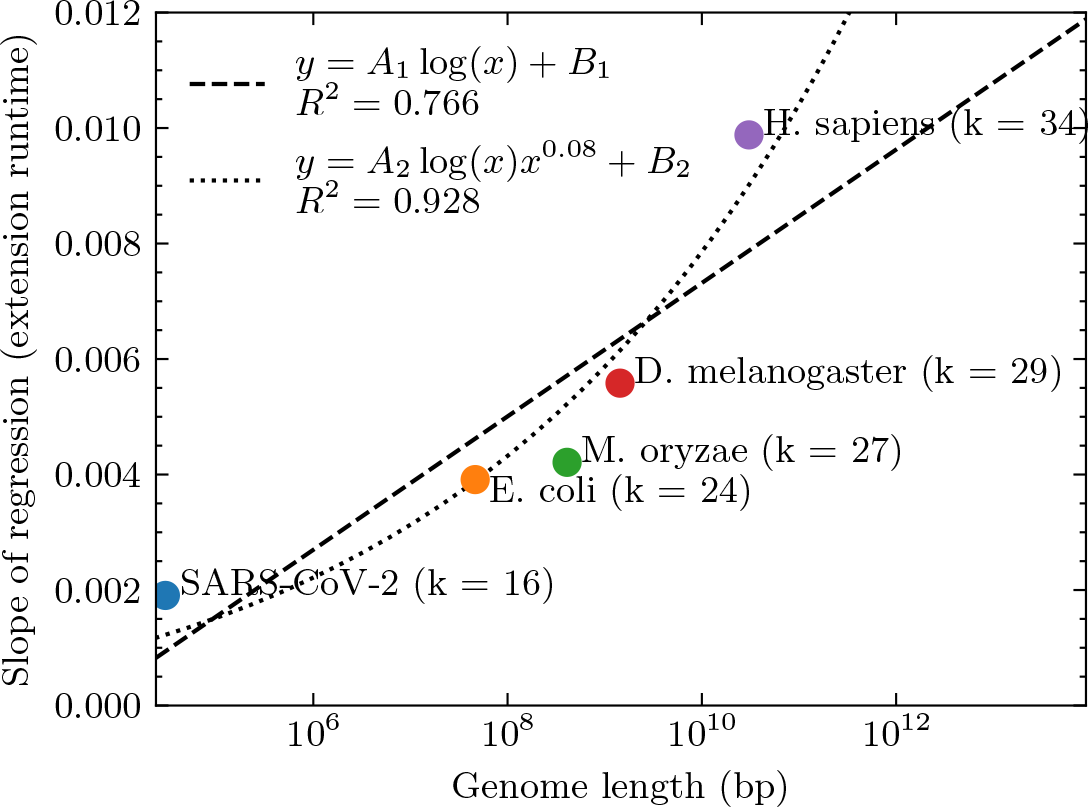
Data points represent the slopes of the linear regressions in Figure 2 for extension, with the value of k (which is *k* = *C* log *n* for a constant 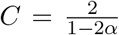) labelled. This dependence of the genome size, *n* (x-axis), is decently approximated by a naive *A*_1_ log(*n*) + *B*_1_ fit where *A*_1_ and *B*_1_ are paraeters. However, our theory states that the dependence should be log(*n*)*n^Cα^* with *Cα* ~ 0.08 when *θ* = 0.05. Fitting *A*_2_ log(*n*)*n*^0.08^ + *B*_2_ gives a better *R*^2^ value (0.928 vs 0.766) with the same number of parameters (2 parameters for both fits), indicating the goodness of our theoretical predictions.

#### Recoverability upper bound by aligned fraction

Recoverability is not measurable on real, non-simulated reads because true sequence homology is not known. However, we can upper bound recoverability by simply measuring the length of the chain relative to the read length, as any homologous bases outside of the chain (i.e. not within the first and last anchor) can not be recovered under our model. We call this the aligned fraction, which our theory also predicts should be > 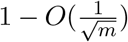 in expectation (with implied constants depending on *θ* and *n*). Practically, the aligned fraction is easily interpretable, and the ability to predict the aligned fraction is useful for ensuring that reads with a given error rate and k-mer size are long enough to align to a reference.

We also observed that the aligned fraction was the main cause of recoverability loss in the simulated experiments (see Supplementary Figure 9), and it turns out that the main 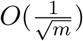 term in the theoretical recoverability result exactly comes from bounding the aligned fraction in the proof (Supplementary Lemma 7), so this quantity is interesting theoretically as well.

In Figure 4, we plot the aligned fraction for each data set. We perform a least-squares fit of the function 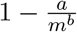 where *a* and *b* are parameters. We exclude points with < 0.25 aligned fraction if the read length is > 1000 when fitting because these points appear to be outliers caused by poor chaining and can bias the least-squares fit. The resulting fits for all plots are reasonable, with *R*^2^ > 0.5 except for the D. melanogaster dataset. The seemingly small value of *R*^2^ can be explained by the fact that most reads (passing our filters) are well-aligned, so the data is almost constant and ≈ 1. Constant data will have *R*^2^ = 0, so we can not do much better in this case. Nevertheless, *b* > 0.85 on all data sets, suggesting that our 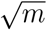 term may be too conservative. However, the exact parameter values in the fit are highly sensitive to the fitting methodology, so the exact values displayed are only suggestive and not to be overinterpreted.

**Fig. 4:**
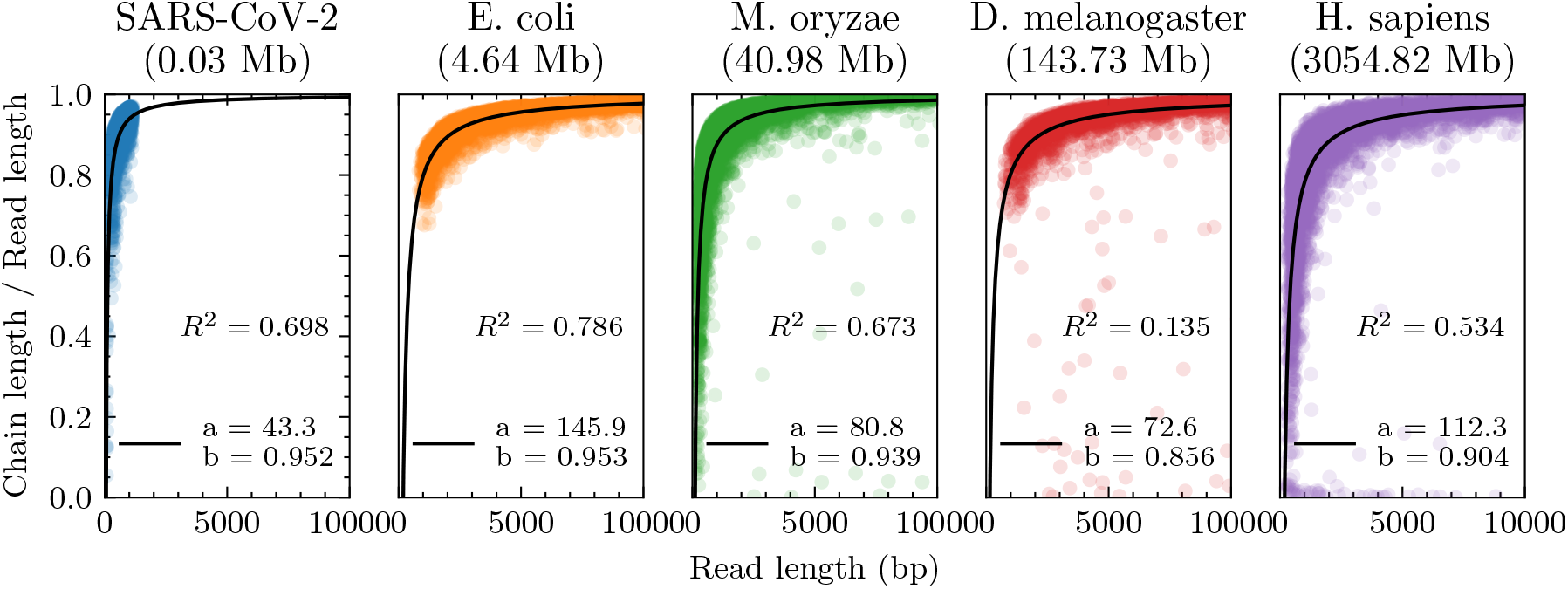
The aligned fraction of the aligned reads, which is chain length divided by read length. We fit a function of the form 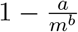 where *m* is the read length (x-axis) and display the resulting *R*^2^ values. Aligned fraction is an upper bound for recoverability, and our upper bound on recoverability (and also aligned fraction) is 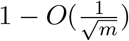, which the data suggests may be too conservative.

**Fig. 5:**
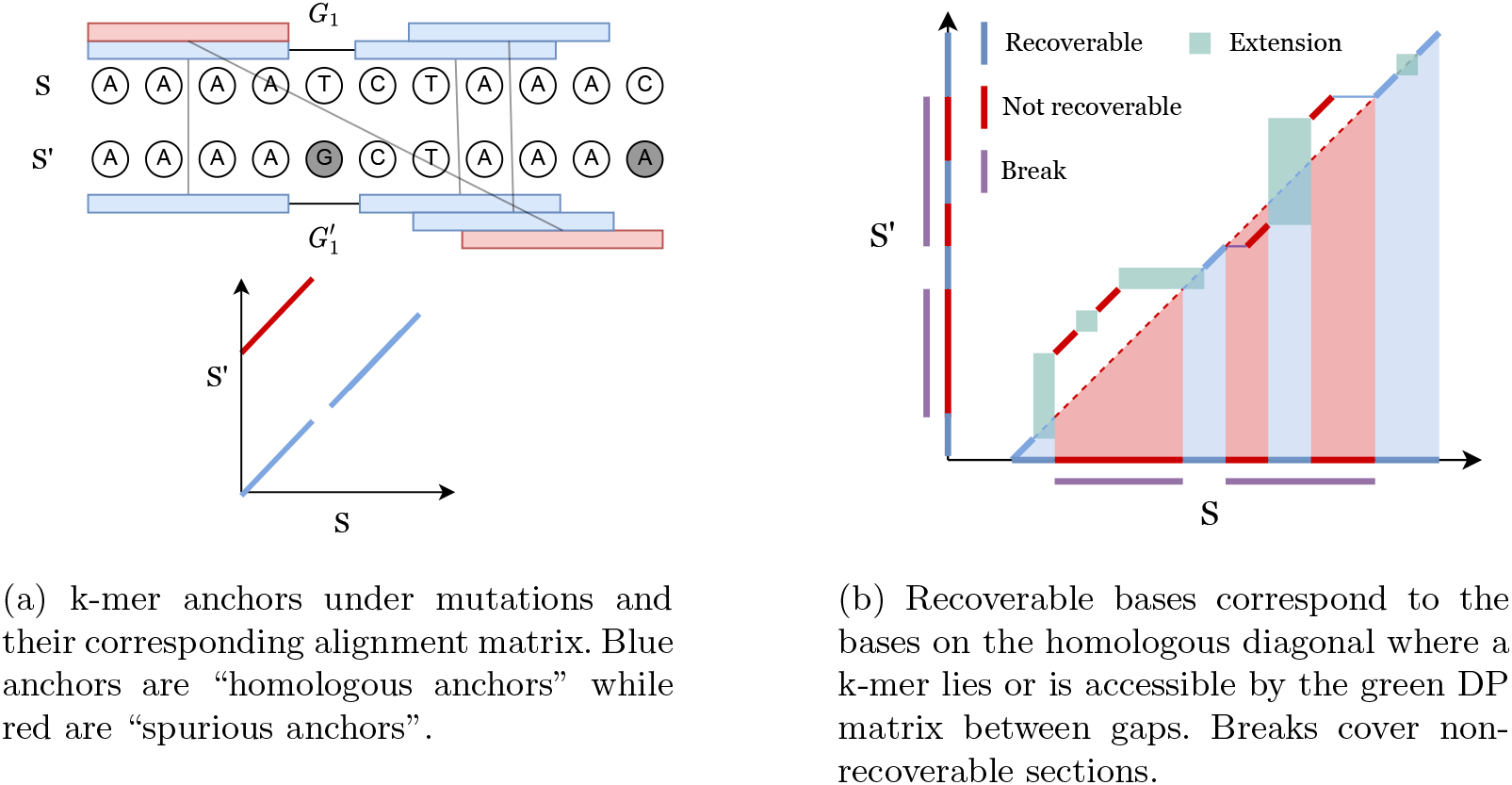
Seed-chain-extend visualized on an alignment matrix. Anchors are short diagonal matches in the matrix. Extension corresponds to performing dynamic programming (DP) on the sub-matrix between anchors. 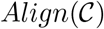 consists of anchors and DP matrices in (b).

## 3 Discussion

In this work, we are able to *rigorously* justify the empirical results seen by the seed-chain-extend heuristic through average case analysis under a simple mutation model. We showed the alignment is both accurate and fast: 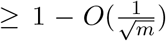 fraction of the sequence homology is recoverable from the chain while only running in *O*(*mn^Cα^* log(*n*)) time, where for even a moderate mutation rate *θ* = 0.05, *Cα* < 0.08. Interestingly, a recent aligner (Koerkamp and Ivanov, 2022) empirically achieved this predicted *n*^1.08^ runtime for *θ* = 0.05 on two length *n* sequences using similar but distinct techniques. In addition, we also proved that one can sketch to arbitrary densities 1/*c* where *c* < *k* = *O*(log *n*) while asymptotically decreasing runtime and without asymptotically decreasing recoverability, justifying the effectiveness of sketching. Because our set-up is modeled by techniques used by practical software such as minimap2, our results provide a theoretical backing for why modern sequence alignment software actually performs so well in practice.

We verified our theoretical results in a synthetic experiment, which showed that our main theoretical predictions were valid for our random model. Specifically, our strongly sub-quadratic bounds predict runtime well, sketching can decrease chaining time without increasing extension time too much, and recoverability is high. Importantly, our results generalize past our uniform string and point substitution assumptions; we show that the runtimes for aligning real nanopore reads to references are still well-approximated by our theory (see Figure 3). However, on more complex, repetitive genomes, our runtime predictions for chaining break down due to real k-mers being much more repetitive than under a uniform random model.

In terms of further work, we propose two possible directions. The first is to optimize the bounds obtained for the current random model proposed in this work, which assumes uniformly random strings and point substitutions. The second is to generalize the random model to model complex genomes more accurately, e.g. repetitive k-mers.

### 3.1 Improving bounds for the current random model

For runtime, it seems unlikely that the expected runtime is truly quasilinear due to the fundamental quantity (1 – *θ*)^*k*^ = (1 – *θ*)^*C* log*n*^ = *n^-Cα^*, the likelihood of a k-mer match, decreasing faster than logarithmically in *n* when *k* = *Θ*(log *n*); this turns out to be the cause of the *n^Cα^* term in the runtime (see Methods). In spite of this, we believe there can be significant improvements for our bounds in Theorems 1 and 2. The most significant aspect which we believe can be improved is the restriction on the constant C. C is required to be 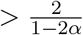, which seems unsatisfactory because this leads to high k-mer sizes, e.g. *k* = 34 for *θ* = 0.05 on the human genome. This is much larger than the k-mer sizes used by noisy long-read aligners in practice (minimap2 defaults to *k* = 15).

The restricted values of C is due to our analysis of spurious anchors relying on a relatively weak variance bound (Lemmas 3 and 4). To use the bound effectively, C must be quite large. The variance bound is also the cause for the 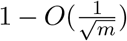 bound for recoverability, which we also believe can be tightened based on both the simulated and real experiments. To surpass these bounds, a deeper understanding of spurious anchor subsequences must be developed. Spurious subsequences of anchors is a very similar problem to common subsequences in random strings for extremely large alphabets, a topic that has had much theoretical attention (Chvátal and Sankoff, 1975; Kiwi et al., 2005; Lember and Matzinger, 2009; Navarro, 2001).

### 3.2 Generalizing the random model

Our current random model does not model complex genomes properly. It is well known that genomes can be extremely repetitive (Koning et al., 2011), which violates the uniform random string assumption. This is why the expected runtime of chaining deviates from our predictions in Figure 2 for the human genome. One idea for modelling repeats is to use a k-mer Markov model for the random strings instead of an independent uniform string. However, the analysis of such models is highly non-trivial (Reinert et al., 2000), and our techniques for analyzing k-mers in this work may not apply. Regardless of how repeats are modelled, any sort of non-trivial analysis would still require a better handling of spurious anchor subsequences, which is exactly the issue that needs to be tackled in order to improve the bounds for our current model.

Another direction is to incorporate indels into the random model. The technical reason we opted for modelling point substitutions instead of indels is that indels complicate the statistics of k-mer matching. For example, an indel of the nucleotide A may not “change” the 4-mers in the sequence AAAAAA, and sequence homology becomes ambiguous. A formulation of an “indel channel” as a model of sequence mutation was used by Ganesh and Sy (2020) for analysis of edit distance, but the resulting analysis for this model was more complex than in the independent substitution case.

Generalizing our model to small indels seems less pressing as our theory was successful for real nanopore data, which has many small indel errors, showing that the substitution model already generalizes well to small indels. However, modelling large structural variations could be an interesting and relevant problem.

## 4 Methods

### 4.1 Homology and recoverability

Let the interval [*a..b*] be the discrete interval {*a, a* + 1,…, *b*} and *S*[*a..b*] be the substring indexed by [*a..b*]. *S′* is a mutated version of the substring *S*[*p* + 1..*p* + *m*] of length *m* starting at *p* + 1, so the optimal alignment should map *S*′ to the homologous indices [*p* + 1..*p* + *m*]. We now define *recoverability* as the number of homologous bases one can possibly recover by seed-chain-extend; a visual example is shown in Figure 5b.

#### Definition 1.

*Let an alignment matrix be given as* [1..|*S*|] × [*p* + 1..*p* + |*S*′|]. *Define the homologous diagonal D_H_* = {(*p* + 1,*p* + 1),…, (*p* + |*S*′|, *p* + |*S*′|)}. *Given a chain* 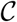, *let* 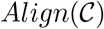 *be the set of possible alignments obtained by extending through gaps and k-mer matching, where* 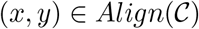 *indicates x and y can be aligned. Then the set of recoverable bases is* 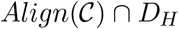 *and* 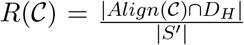.

We define 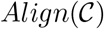 carefully in the appendix. Note that an optimal chain with respect to our linear gap cost objective *u* – *ζ*[(*i_u_* – *i*_1_) + (*j_u_* – *j*_1_)] does not directly optimize for recoverability. We provably find the chain with the optimal linear gap cost, and then we will argue that it leads to good recoverability.

Traditionally, alignment optimality is defined using a generalized edit distance (Vingron and Waterman, 1994). However, these distances are only used as a proxy for detecting sequence homology (Batzoglou, 2005; Thorne et al., 1991; Lunter et al., 2005). We know the true underlying sequence ancestry in our model, so defining optimality with respect to sequence homology suits the actual goal of sequence alignment. The reason for the name “recoverability” is that extension could *potentially* recover all recoverable bases, but this depends on the extension algorithm and is not guaranteed (Lunter et al., 2008).

Under our model, the trivial O(1) alignment that aligns *S*′ back to the originating substring of *S* without indels is the most homologous. This may seem to make our results superfluous; however, remember that the algorithm does not know where *S*′ “begins” if *m* < *n*. Also, while we do not attempt the case with indels, seed-chain-extend is still valid when indels are present whereas the trivial solution does not allow for indels.

We will lower bound 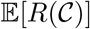. To do this, we will work with a more natural object called a *break*.

#### Definition 2 (inspired by Ganesh and Sy (2020)).

*We call matching bases and anchors of the form* (*x,x*) *homologous and spurious otherwise. Given a chain* ((*i*_1_, *j*_1_),…, (*i_u_, j_u_*)) *and a maximal interval* [*p..q*] *such that* (*i_p_, j_p_*),…, (*i_q_, j_q_*) *are all spurious anchors, define the break B as B* = [min(*i_p_, j_p_*).. max(*i_q_, j_q_*) + *k* – 1]. *Let the length or size of a break be L*(*B*) = max(*i_q_, j_q_*) – min(*i_p_, j_p_*) + *k and* 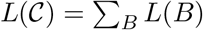 *be the total length over all breaks*.

#### Lemma 1.

*Given any chain* 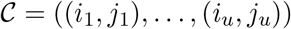, *we have that* 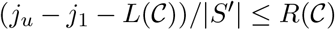.

We prove Lemma 1 in the appendix. The concept of a break is illustrated in Figure 5b. Breaks cover non-recoverable regions, so subtracting the breaks from the span of the anchors lower bounds the recoverability.

### 4.2 Fundamental tools and bounds

In this section, we describe some fundamental tools for dealing with pairs of random mutating strings. We first need to give a careful probabilistic exploration of random k-mer anchors on *S* = *x*_1_ *x*_2_ … and *S*′ = *y_p_*+1*y*_*p*+2_ …. This requires a bit of work due to the dependence between the random strings *S, S*′. For the rest of the paper, missing proofs can be found in the appendix.

#### Definition 3.

*Let M*(*i, j*) *be random variables such that M*(*i, j*) = 1 *if x_i_* = *y_j_ and* 0 *otherwise. Let A*(*i, j*) *be a random variable* 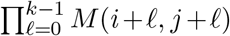. *A*(*i, j*) *is an indicator random variable for the presence of a k-mer anchor at positions* (*i, j*). *We will also refer to A*(*i, j*) *variables as “anchors”*.

We would like for *M*(*i, j*), *M*(*i*+1, *j*+1),… to be independent, so finding the probability of k-mer matches is easy. However, in our model, it is not actually obvious *a priori* that *M*(*i, j*), *M*(*i*+1, *j* + 1) are independent when *j* ≠ *i*. Consider *M*(1, 2) and *M*(2,3). The former is a function of *x*_1_, *y*_2_, and the latter *x*_2_, *y*_3_. However, *x*_2_ and *y*_2_ are dependent in our model, so more work is needed. A convenient graphical representation of independence for the *M* random variables is the *match graph*, which we define below.

#### Definition 4.

*The match graph of a set of random variables of the form* 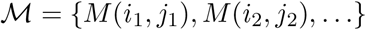 *is a graph* 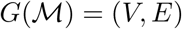 *where the vertices V are the letters x*_1_,…, *x*_*n*+*k*-1_, *y*_*p*+1_,…, *y*_*p*+*m*+*k*–1_, *and the edges*

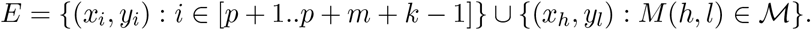

*In particular, the match graph is bipartite for the sets* {*x*_1_,…, *x*_*n*+*k*-1_} *and* {*y*_*p*+1_,…, *y*_*p*+*m*+*k*–1_}.

A match graph is shown in Supplementary Figure 1. The main reason for defining the match graph is the following theorem, which allows us to graphically determine independence of the *A* variables.

#### Theorem 3.

*The random variables* 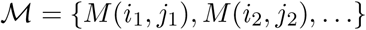 *where i*_ℓ_ ≠ *j_ℓ_ for all* 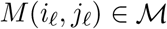 *are independent if the induced match graph has no cycles*.

#### Corollary 1.

*If i* ≠ *j*, 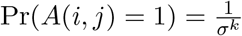. *Otherwise*, Pr(*A*(*i,i*) = 1) = (1 – *θ*)^*k*^.

We will denote the random variables 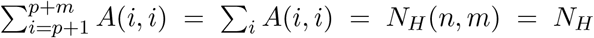 and 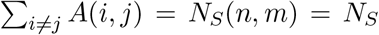 respectively as the number of homologous anchors and spurious anchors; we will drop the dependence on *n, m* to simplify notation. These are key random variables that we wish to bound later on. The below theorem follows directly from Corollary 1 by linearity of expectation.

#### Theorem 4.

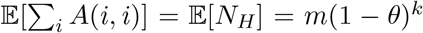, *and* 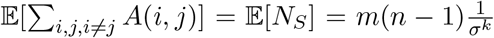. *In particular*, 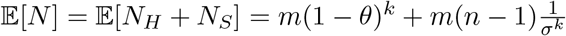.

### 4.3 Bounding sums of k-mer random variables

We proceed to bound the *distribution* of the random variables *N_S_* and *N_H_*, which are sums of *dependent* random variables. We first bound *N_S_* by computing the second moments and using variance based bounds. To do this, we need to examine the independence structure of the *A*(*i, j*) random variables.

#### Lemma 2.

*For A*(*i,j*) *and A*(*h, l*), *if both of the following conditions hold*:

1. |*i* – *h*| ≥ *k or* |*j* – *l*| ≥ *k and*
2. |*i* – *l*| ≥ *k or* |*j* – *h*| ≥ *k*, *then the induced match graph on the M variables for A*(*i,j*) *and A*(*h, l*) *has no cycles*.

Intuitively, the first condition states that two anchors do not overlap too much, e.g. *A*(1,1) and *A*(2, 2) are not independent when *k* = 3. The intuition behind the second condition can be illustrated by the following situation where *k* = 1 and *θ* ~ 0: consider the anchors *A*(1, 5), *A*(5, 1). Since it is likely that *x*_1_ = *y*_1_ and *x*_5_ = *y*_5_, if *x*_1_ = *y*_5_ then *x*_5_ = *y*_1_ with high probability, so *x*_1_ = *y*_5_ is not independent of *x*_5_ = *y*_1_.

#### Corollary 2.

*If A*(*i,j*) *and A*(*h, l*) *satisfy Lemma 2, they are independent*.

*Proof.* By condition (1) in Lemma 2, *A*(*i,j*) and *A*(*h, l*) do not share any *M* variables i.e. *A*(*i, j*) = *M*(*i, j*)*M*(*i* + 1, *j* + 1)… and similarly for *A*(*h, l*), but no product shares a variable with the others. Since the match graph has no cycles under these conditions, by Theorem 3, all *M* variables are independent so 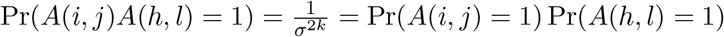 as desired.

#### Lemma 3.

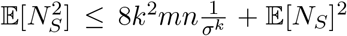. *Thus the variance Var*(*N_S_*) *can be upper bounded by* 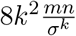. *Furthermore*, 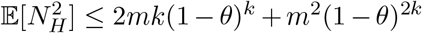 *and* 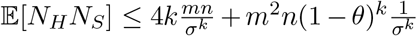.

Now we can use the variance bound and Chebyshev’s inequality to get the result below. Note the bound uses *k* = *C* log *n*; we will prefer this form for quantities directly used for proving the main result. The label F1 in the theorem refers to the particular event space for which the bound always holds. We will label each proposition that holds with high probability with the event space that we are operating in. We will continue this convention for the rest of the paper as this will be useful when computing our final bounds.

#### Lemma 4 (F1).

*With probability* ≥ 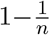, *the number of spurious anchors is* 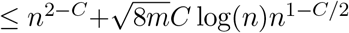 *where we used* 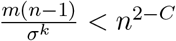. *That is*,

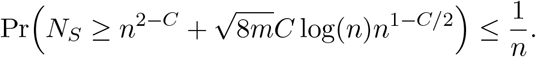

If *C* > 3, then for large *n, N_S_* = 0 with high probability, and our analysis would be easy. But, we want C as small as possible. It turns out we can make *C* ~ 2 for reasonable *θ*, significantly tightening our bounds.

For *N_H_* we can get a stronger exponential bound because of its independence structure. *A*(*i, i*)s, which we call homologous anchors, are only dependent in a small neighbourhood around *i* of size *k* because k-mers on non-overlapping substrings are independent. This is called *k*–*dependence* (not to be confused with *k-independence*) and is used in (Blanca et al., 2022) to show *N_H_* is asymptotically normal. Concentration bounds can also be translated in the *k*–dependent scenario (Janson, 2004).

#### Theorem 5 (Dependent Chernoff-Hoeffding bound - Corollary 2.4 from Janson (Janson, 2004) reworded and simplified).

*Suppose we have* 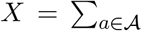 *Bernoulli_a_*(*q*) *for some* 0 < *q* < 1. *A proper cover of* 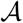 *is a family of subsets* 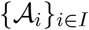 *such that all random variables in* 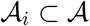 *are independent and* 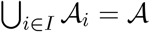. *Let* 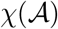 *be the minimum size of the cover*, |*I*|, *over all possible proper covers. Then for t* ≥ 0,

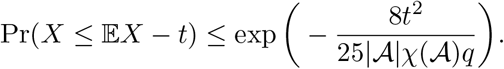

#### Lemma 5.

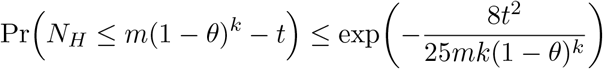

*Proof.* We simply use Theorem 5 with *q* = (1 – *θ*)^*k*^. By k-dependence, we can easily see that 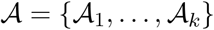 where 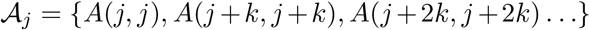 is a partition satisfying the independence condition, and we will have at most *k* sets. Thus 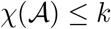, and we’re done.

### 4.4 Proof of non-sketched main result

To prove the main result on seed-chain-extend without sketching, we will need to bound three quantities in expectation: the runtime of chaining, the recoverability of chaining, and the runtime of extension.

We first bound 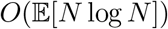, the expected runtime of chaining (and also anchor sorting). Note that 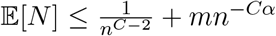. We would like 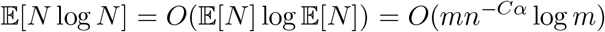 to hold. Unfortunately, Jensen’s inequality only gives 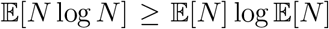 because *x* log *x* is convex. However, with a bit more work:

#### Theorem 6.

*Assume m* = *Ω*(*n*^2*Cα+ε*^) *for some ϵ* > 0, *and* 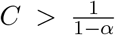. *Letting N be the total number of k-mer anchors*, 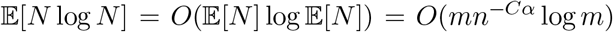. *Thus the runtime of chaining is O*(*mn^-Cα^* log *m*).

Now we bound the expected recoverability of the chain. Given *S* and *S*′, let a *homologous gap* of size ℓ + *k* – 1 bases be an interval of ℓ consecutive k-mers for which no homologous anchors exist (i.e. the k-mers are mutated). In the context of a chain, a homologous gap will refer to a gap flanked by two homologous anchors. Technically, if ℓ consecutive k-mers are uncovered but are flanked by two homologous anchors, this gives ℓ – *k* + 1 uncovered bases. We will ignore these factors of k as we will show that they are asymptotically small. It turns out homologous gaps grow relatively slowly in *n* with high probability.

#### Lemma 6 (F2).

*With probability* ≥ 1 – 1/*n, no homologous gap has size greater than*

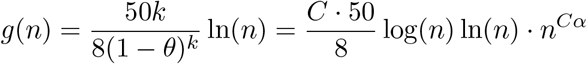

*plus a small C* log *n term we will ignore because it is small asymptotically*.

In a chain, gaps may also be flanked by one or two spurious anchors. We call these *non-homologous gaps*. We first bound break lengths, which will imply good recoverability, and bound non-homologous gaps later on.

#### Lemma 7 (F1 + F2).

*Take any* 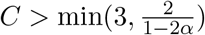 *and let* 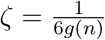 *where* 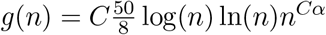 ln(*n*)*n^Cα^. Assume m* = *Ω*(*n*^2*Cα+ϵ*^) *for some ϵ* > 0. *Then for large enough n, there are no breaks of length* ≥ *m*^1/2^ *with probability greater than* 1 – 2/*n in an optimal chain*.

#### Corollary 3.

*Under the same assumptions as in Lemma 7, the expected recoverability of any optimal chain is* 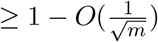.

The idea behind proving the above propositions is to work in a space of events F1 ⋂ F2 where “bad events” do not occur and any optimal chain has good recoverability. Because this space of bad events is small, they do not contribute to our expected value too much. This finishes the recoverability result for the main theorem.

The last step is to bound extension running time, and this comes down to separately bounding the size of the homologous and non-homologous gaps in any optimal chain. We bound the runtime of extension through homologous gaps by directly calculating the expectation through *all possible* homologous gaps. We then show that the runtime through non-homologous gaps is small and does not contribute to the asymptotic term.

#### Lemma 8.

*Let* 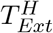 *be the time of extension over only the homologous gaps of any optimal chain*. 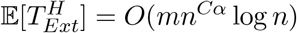.

#### Lemma 9.

*Let* 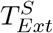 *be the runtime of extension through only the non-homologous gaps of an optimal chain. Under the same assumptions as in Lemma 7*, 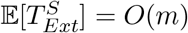.

*Proof (Theorem 1).* The expected runtime of chaining follows from Theorem 6. The recoverability result follows from Corollary 3. The expected runtime of extension is 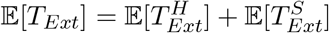, and 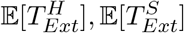 are both *O*(*mn^Cα^* log(*n*)) by Lemmas 8 and 9. This completes the proof as long as we satisfy the assumptions of Lemma 7 and Theorem 6 on *C, α*. To satisfy the assumptions, we require 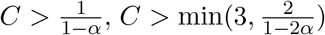, and *Cα* < 1/2 otherwise *m* = *Ω*(*n*^2*Cα*+*ε*^) > *n* for large enough *n*. It’s not hard to check that the limiting condition is *α* < 1/6, so we require – log_4_(1 – *θ*) < 1/6. This works out to be 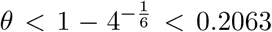. We can also remove the minimum condition on *C* because *α* < 1/6 implies 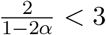.

### 4.5 Sketching and local k-mer selection

Now consider not selecting all of the k-mers in a string, but only a subset of them during the initial seeding step. This allows one to chain only a subset of the k-mers, potentially providing runtime savings.

We use the *open syncmer* method (Edgar, 2021). Given a string, we take all k-mers of the string and break the k-mer into s-mers with *s* < *k*. There are *k* – *s* + 1 s-mers in the k-mer. We select or *seed* the k-mer if the smallest s-mer (subject to some ordering, which we choose as uniform random) is in the 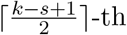 1-indexed position; we call a selected k-mer an *open syncmer*. Given repeated smallest s-mers, we take the rightmost one to be the smallest. Finding the smallest s-mer among the *k* – *s* + 1 s-mers in a k-mer takes *k* – *s* + 1 iterations, so finding all open syncmer seeds in *S*′ takes *O*((*k* – *s* + 1)*m*) = *O*(*mk*) = *O*(*m* log*n*) time.

The expected fraction of selected k-mers over a string with i.i.d uniform letters is called the *density*, and it is 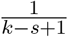 for the open syncmer method (up to a small error term 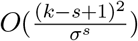which we will ignore; see (Zheng et al., 2020) or (Shaw and Yu, 2022)). We will let c be the reciprocal of the density, so *c* = (*k* – *s* + 1).

The original open syncmer definition in (Edgar, 2021) had a parameter *t* where a k-mer was selected if the smallest s-mer was in the *t*–th position; we proved in (Shaw and Yu, 2022) that the optimal *t* is 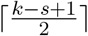 with respect to maximizing the number of conserved bases from k-mer matching. The reason we choose open syncmers is primarily to the following fact which was shown in (Edgar, 2021):

#### Theorem 7.

*Define* 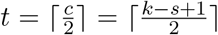. *If k* – *s* + 1 *is odd, two consecutive open syncmers must have starting positions* ≥ *t bases apart. If even, they must have starting positions* ≥ *t* – 1 *bases apart*.

Theorem 7 follows by examining the smallest s-mer in a k-mer and noticing that in the next overlapping k-mer, the locations for the new smallest s-mer are restricted. This theorem is the reason we use open syncmers and is crucial to our proofs. The spacing property makes selected open syncmers a polar set (Zheng et al., 2021); other methods also give rise to polar sets (Frith et al., 2020, 2022) but open syncmers seem to perform well empirically (Shaw and Yu, 2022; Frith et al., 2022; Dutta et al., 2022) and are easy to describe. For the rest of the section, we will assume *c* = *k* – *s* + 1 is odd, so 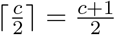.

Let *A*(*i, j*) be the random variables as defined before. Let *J*(*i*) be the indicator random variable for if the *i*th k-mer is selected on *S* as an open syncmer, and *J*′(*j*) similarly for the *j*-th k-mer on *S*′. We now wish to calculate 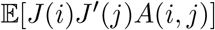, the probability that a k-mer match exists and the k-mer is an open syncmer.

#### Lemma 10.

*If i* = *j, then* 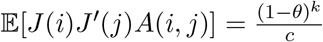. *Otherwise*, 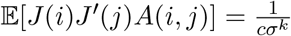.

*Proof.* If *A*(*i, j*) = 1, then the *i*-th k-mer and *j*–th k-mer are the same. If a k-mer is selected as an open syncmer on *S*, it must also be selected on *S*′, so 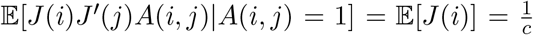. *A*(*i, j*) and *J*(*i*) are independent because we assume the random ordering for the s-mers is independent of the random mutations. Using the law of total expectation and Theorem 4 gives the result for both cases.

#### Definition 5.

*We will replace all random variables involving anchors and matches with a superscript* * *to indicate sketched seeds, e.g. A*(*i,j*)* = *A*(*i,j*)*J*(*i*)*J*′(*j*), 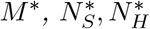, *etc*.

#### Corollary 4.

*The expected total number of anchors after applying open syncmer seeds with density* 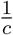 *is*

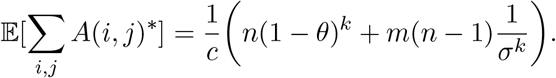

The above follows directly from Lemma 10. As expected, subsampling to a 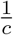 fraction of the k-mers gives 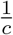 expected hits. Note the important property of *context independence* used in the proofs: if *A*(*i, j*) = 1, then *J*(*i*) = *J*′(*j*). This property is not satisfied if one were sampling k-mers randomly or using minimizers (Roberts et al., 2004; Shaw and Yu, 2022). We now deduce the sketched moment bounds on *N*\*.

#### Lemma 11.

*The variance* 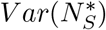 *can be upper bounded by* 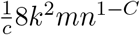. *Furthermore*, 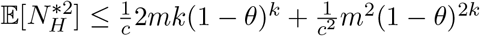 *and* 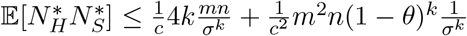.

#### Lemma 12 (F1*).

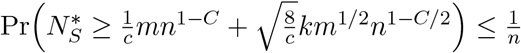

The sketched moment bounds allow us to start re-analyzing the crucial propositions in Section 4.4 in the context of sketching. The first main result is that sketching reduces the chaining time as one would expect.

#### Theorem 8.

*Under the same assumptions as in Theorem 6, letting N*\* *be the total number of sketched k-mer anchors, the expected chaining time is* 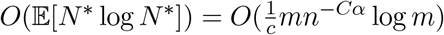.

The second result we wish to highlight is the new bound on extension runtime through homologous gaps.

#### Lemma 13.

*The expected runtime of sketched extension through the homologous gaps in an optimal chain is* 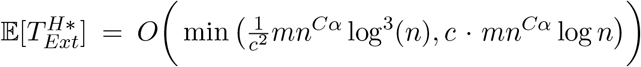. *If c* = *Θ*(log*n*) *then* 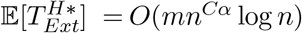.

It will turn out that in the sketched case, extension over homologous gaps also dominates runtime. Since we sketch with density 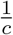, this result states that we can sketch with *decreasing density* and have the *same* asymptotic extension time as without sketching, which is surprising. The crucial fact we use in the proof of Lemma 13 is Theorem 7, which shows that open syncmer seeding weakens the k-dependence of the seeds since they are now at least (*c*+1)/2 bases apart. This tightens the bound in Theorem 5 and also shows that the sketched maximum homologous gap size is *Θ*(*g*(*n*)), with *g*(*n*) as in Lemma 6. If we used other methods such as closed syncmers (Edgar, 2021) or FracMinHash (Irber et al., 2022; Hera et al., 2022), then we would recover the *O*(*c* · *mn^Cα^* log*n*) result in Lemma 13 but not the 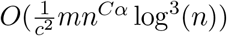 result. Thus, we have saved a log factor by using open syncmers when we let *c* = *Θ*(log*n*). The bulk of Section E in the appendix is dedicated to proving Lemma 13.

The key quantities used for the rest of the proofs are 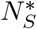 and *g*′(*n*), the new maximum homologous gap size. We will prove *g*′(*n*) = *Θ*(*g*(*n*)) with *g*(*n*) as in Lemma 6, so the only asymptotic difference in these bounds is a factor of 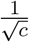 in 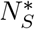. Since we want 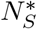 to be small, this does not affect downstream analysis. It follows that the proofs of the rest of the lemmas in Section 4.4 can be replicated almost verbatim. We give a sketch of these proofs in the appendix.

## 5 Software availability

Scripts for regenerating the simulated experiments are available at https://github.com/bluenote-1577/basic_seed_chainer/. The aligner used for the real nanopore experiments is available at https://github.com/bluenote-1577/sce-aligner/ along with scripts for regenerating figures.

## 6 Competing interest statement

The authors declare no competing interests.

## 7 Acknowledgements

J.S. was supported by an NSERC CGS-D scholarship. This work was supported by Natural Sciences and Engineering Research Council of Canada (NSERC) grant RGPIN-2022-03074.

## A Mathematical conventions and reminders

We list a few reminders and conventions that will be useful for the technical parts of our work. ≫ will mean asymptotically dominates, i.e. *f*(*n*) ≫ *g*(*n*) if and only if lim_*n*→∞_ *g*(*n*)/*f*(*n*) = 0. log(*n*) = log_*σ*_(*n*) and ln(*n*) uses the *e* as the base, where *σ* is the alphabet size. |*S*| ~ *n* and |*S*′| ~ *m*, where *S, S*′ are our pairs of strings and ~ ignores small factors of *k* present. This is because *k* = *C* log *n* for a constant *C* > 0 and *m* = *Ω*(*n*^2*Cα*+*ϵ*^) for the assumptions of our main theorems, and these terms dominate *k* = *O*(log *n*). *S*′ is a mutated substring of *S*, where the substring starts at position *p*. (1 – *θ*)^*k*^ = *n^-Cα^* where *α* = – log_*σ*_(1 – *θ*) > 0 is a function of 0 < *θ* < 1, the mutation rate, and *σ* is our alphabet size which is > 1. Our results are general, but we will use *σ* = 4 for specific numerical results. When we use the assumption 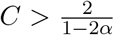, *C* is also greater than 2, a useful fact we will use repeatedly.

A chain 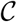 is a sequence of anchors (exact matches of k-mers), represented by tuples ((*i*_1_, *j*_1_),…, (*i_u_, j_u_*)) where *i*_ℓ_ is the starting position of the ℓth k-mer on *S* and *j*_ℓ_ is the starting position of the ℓth k-mer on *S*′. Anchors can overlap. The cost for a chain 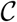 is *u* – *ζ*[(*i_u_* – *i*_1_) + (*j_u_* – *j*_1_)] where *ζ* = *ζ*(*n*) > 0 will eventually be considered as a (decreasing) function of *n*.

## B Missing proofs and definitions from “Homology and recoverability”

We note that technically, the alignment matrix in Definition 1 is slightly different than the standard dynamic programming matrix (Durbin et al., 1998) which can be thought of a directed graph with a path representing an alignment. Our representation does not include the graphical information; it only captures information pertaining to the possibility of matching bases, which is sufficient for us.

### B.1 Recoverability and definition of 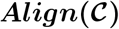

Figure 5b will serve as a helpful guide for our definitions. Let 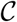 be a chain of anchors ((*i*_1_, *j*_1_),…, (*i_u_, j_u_*)). Below we define carefully define recoverability and set up our proof of Lemma 1.

A chain gives a set of k-mer matches and a set of possible alignments by extending through bases, constraining the full alignment matrix between *S* and *S*′. We can formalize this as follows. Given two consecutive anchors (*i_ℓ_, j_ℓ_*) and (*i*_ℓ+1_, *j*_ℓ+1_), extending allows for possible matches between the gaps given by the following set of possible matches

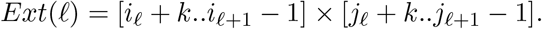

The factor of +*k* and –1 is because *i_ℓ_* is the start of the ℓth anchor, and extension starts after the end of the first k-mer and goes until 1 base before the start of the second k-mer. *Ext*(ℓ) corresponds to the green boxes in Figure 5b. The set *Ext*(ℓ) is empty if *i_ℓ_* + *k* > *i*_ℓ+1_ – 1 and similarly for *j*, i.e. there are no gaps between possibly overlapping anchors. 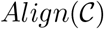 also takes into account the k-mer matches given by 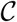. Thus we define 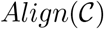 formally as

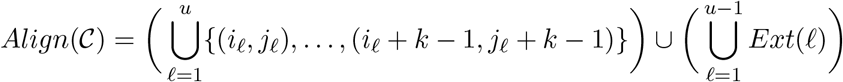

where the first term corresponds to the k-mer matches in the alignment matrix.

### B.2 Proof of Lemma 1

Since we care about the intersection of 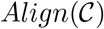 with the homologous diagonal, we can manipulate the expression for 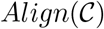 to get something more tractable. Let Diag[*a..b*] = {(*x,x*) : *x* ∈ [*a..b*]}, 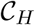 be the set of homologous anchors in 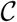, and 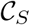 be the set of spurious anchors. The homologous anchors give rise to the homologous matches in the alignment matrix

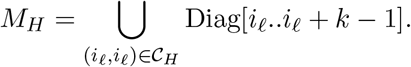

The spurious anchors give no matches on the homologous diagonal, so it contributes nothing to the 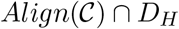 term. Therefore,

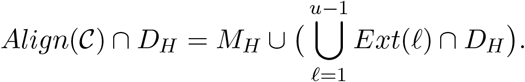

We define

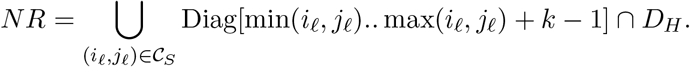

NR represents the parts of the diagonal in Figure 5b that are not recoverable or accessible by extension through anchors (although it may technically intersect 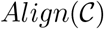 in our mathematical definition). The following identity then holds after rewriting *M_H_*:

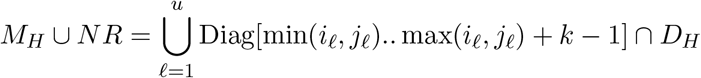

because the min-max condition is redundant for homologous anchors where *i_ℓ_* = *j_ℓ_*. We can fill in the gaps between *M_H_* ∪ *NR* along the diagonal with *Ext*(ℓ) after noticing that we can rewrite

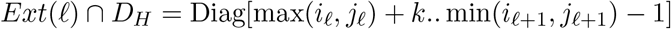

using the fact that (*x, x*) ∈ *Ext*(ℓ) ⇔ *i_ℓ_* + *k* ≤ *x* ≤ *i*_ℓ+1_ – 1 and *j_ℓ_* + *k* ≤ *x* ≤ *j*_ℓ+1_ – 1. Finally,

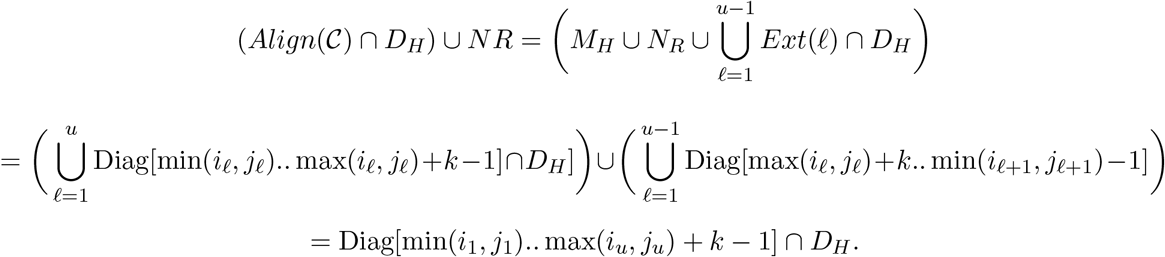

This result follows because the union over all sets covers the entire diagonal between the first and last anchor. After bounding the diagonal to lie within *D_H_* and removing *NR* from both sides of the equation, we obtain the following result.

#### Supplementary Lemma 1.

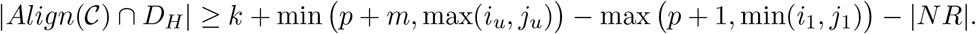

*Proof.* The following inequality holds by simple set theoretic arguments:

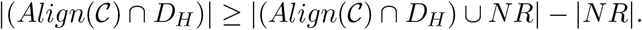

Now notice that the term Diag[min(*i*_1_, *j*_1_).. max(*i_u_, j_u_*) + *k* – 1] may technically lie outside the diagonal [*p* + 1..*m*] × [*p* + 1..*m*], so to constrain it to lie on the diagonal after intersecting with *D_H_*, we must make max(*i_u_, j_u_*) = *p* + *m* if *i_u_* > *p* + *m* and min(*i*_1_, *j*_1_) = *p* + 1 if *i*_1_ < *p* + 1. Thus the cardinality of the set Diag[min(*i*_1_, *j*_1_).. max(*i_u_, j_u_*) + *k* – 1] ⋂ *D_H_* is *k* + min(*p* + *m*, max(*i_u_, j_u_*)) – max(*p* + 1, min(*i*_1_, *j*_1_)), which finishes the proof.

#### Lemma 1

*Given any chain* 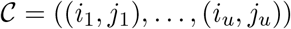, *we have that* 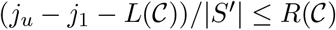.

*Proof.* Using the supplementary lemma above, we note that *j_u_* ≤ *p* + *m* and *j*_1_ ≥ *p* + 1 since the anchors must lie between [*p* + 1..*p* + *m*], so after ignoring the *k* term we get

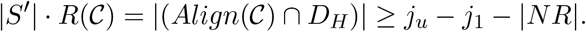

We now prove that the breaks “cover” *NR*, so |*NR*| ≤ *L* where *L* is the total length of the breaks, proving the result. Let *π*(*NR*) be the projection of the set onto one of the coordinate axes (it doesn’t matter which one). Let *x* ∈ *π*(*NR*). Then for some spurious anchor (*i, j*), min(*i, j*) ≤ *x* ≤ max(*i, j*) + *k* – 1. But every spurious anchor is contained in exactly one break, since breaks partition the set of spurious anchors. This break *B* = [a..b + *k* – 1] ⊃ [min(*i, j*).. max(*i, j*) + *k* – 1] since the break takes the minimum and maximum coordinates over *all* spurious anchors in the break. Thus *x* ∈ *B* for some break *B* and the set of breaks 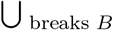, in 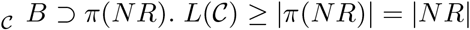, and we’re done.

## C Missing proofs from “Fundamental tools and bounds” and “Bounding sums of k-mer random variables”

### C.1 Proof of Theorem 3

**Supplementary Figure 1:**
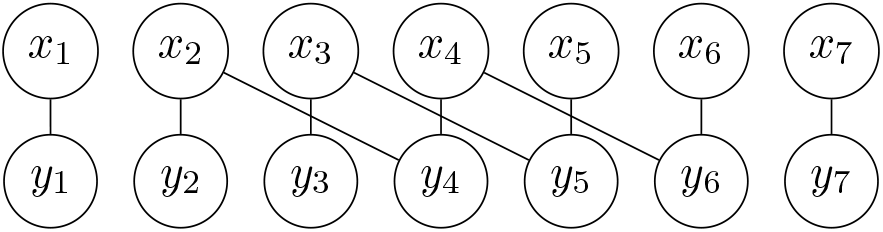
A match graph 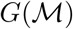 where 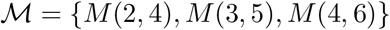 and |*S*′| = |*S*|.

Intuitively, a match graph encodes dependencies between random letters after conditioning on 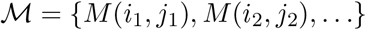. The main idea is that for such a graph, the connected components should be mutually independent of everything not in the component because all of the dependencies lie only in that connected component. The match graph is an example of a *dependency graph*; see Chapter 5 in (Alon and Spencer, 2015).

#### Supplementary Lemma 2.

*Consider a set of random variables of the form* 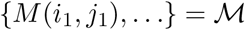 *If two vertices x,y are in separate connected components in* 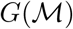 *then they are conditionally independent of* 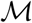.

*Proof.* Consider the connected components *G*_1_,…, *G_q_* of 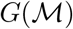) and the edges within the connected components corresponding to the random variables in 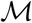; call this partition of edges {*E*_1_,…, *E_q_*} where 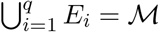.

First, we claim that *G*_1_,…, *G_q_*, considered as random letters in *S* and *S*′, are mutually independent (without considering additional 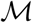 variables). Indeed, each *G_i_* can be considered as just a set of pairs of (*x_ℓ_, y_ℓ_*) random variables because the edges (*x_ℓ_, y_ℓ_*) always exist in the original graph and so the connected component must leave no *x_i_* or *y_i_* unpaired. All pairs of (*x_ℓ_, y_ℓ_*) random variables are mutually independent by definition of our original mutation model. Thus all *G_i_*s are mutually independent.

Now we consider the *E_ℓ_*s. The random variables in *E_ℓ_* are functions of *G_ℓ_*, the letters in the connected component. It follows that the *E_ℓ_*s are functions of mutually independent *G_ℓ_*s, and are themselves mutually independent. Now let *x* ∈ *G_x_* and *y* ∈ *G_y_* be two letters in different connected components. *x, E_x_* are both functions of *G_x_* and similarly for *y, E_y_*, and *G_y_*. Thus 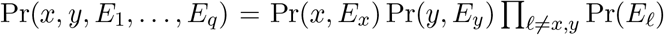 which shows conditional independence of *x, y* with respect to 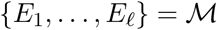.

#### Theorem 3

*The random variables* 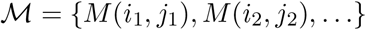 *where i_ℓ_* ≠ *j_ℓ_ for all* 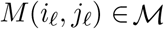 *are independent if the induced match graph has no cycles*.

*Proof.* We shorten *M*(*i_ℓ_, j_ℓ_* = *M_ℓ_*. We will proceed by induction on the size of 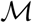. The case 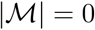 is vacuously true. If 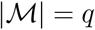, We want to show that Pr(*M*_1_,…, *M_q_*) = Pr(*M*_1_,…, *M*_*q*–1_) Pr(*M_q_*); the first term is 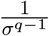 because *M*_1_,…, *M*_*q*–1_ must also be cycle free and by the induction assumption. Let 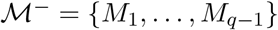. Let *M_q_* = *M*(*α, β*). We need to calculate 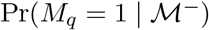 and show that it is equal to 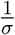.

By definition, we have that

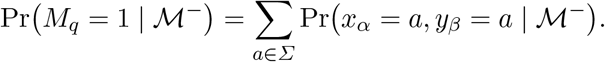

Since 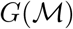 has no cycles, *x_α_* and *y_β_* lie on two different connected components of 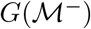 after removing the edge induced by *M*(*α, β*) (we use the *α* ≠ *β* assumption here). Therefore *x_α_, y_β_* are conditionally independent by the above Supplementary Lemma 2, so we can write the above sum as

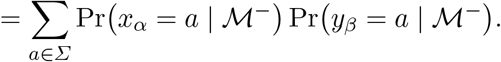

We claim that both terms in the sum are 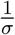. To show this, let the permutation *C_σ_* be a cyclic permutation of order *σ* on letters in *Σ* (e.g. *C*_4_ sends *A* → *C* → *T* → *G*) and apply it to every letter on both strings when we write *C_σ_*(*S, S*′). It’s not hard to see that this is a measure (or probability) preserving transformation for 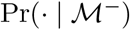 because it preserves matches between *S* and *S*′ and letters are distributed uniformly.

Let *A* be the set of strings (*S, S*′) for which *x_α_* = *a*, and we can see that 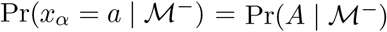. Now applying *C_σ_ σ* times and using measure preservation gets that

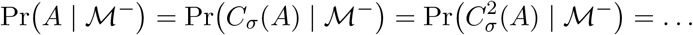

Finally, all 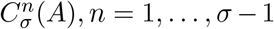 are clearly disjoint and partitions the space of strings. Thus, we see that 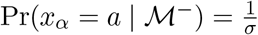 as desired. The argument works exactly the same for 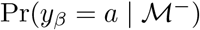, so

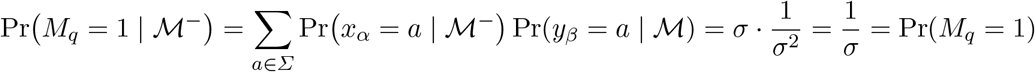

and our induction step is complete.

### C.2 Proof of Corollary 1

#### Corollary 1

*If i* ≠ *j*, 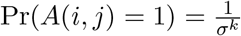. *Otherwise*, Pr(*A*(*i, i*) = 1) = (1 – *θ*)^*k*^.

*Proof.* If the match graph induced by *A*(*i, j*), i.e. induced by *M*(*i, j*), *M*(*i* + 1, *j* + 1),…, *M*(*i* + *k* – 1, *j* + *k* – 1) has no cycles, using Theorem 3 gives the result. This is easy to see from drawing out the match graph which has edges (*x, y*) for all *x, y* and (*x_i_, y_i_*),…,(*x*_*i*+*k*-1_, *y*_*j*+*k*-1_) but we give a rigorous argument below. See Supplementary Figure 1 for an example of such a match graph.

First, suppose *j* > i and suppose *x*_*i*+1_ is the vertex with a cycle and with the smallest possible ℓ ∈ [0..*k* – 1]. We can assume the cycle touches some x because the match graph is bipartite. Since *x*_*i*+1_ has degree at most two and therefore equal to two, we can traverse the edge (*x*_*i*+ℓ_, *y*_*i*+ℓ_) as the first edge in the cycle. If *y*_*i*+1_ has degree one, then *x*_*i*+1_ has no cycle so suppose the degree is two. Then we have an edge (*y*_*i*+ℓ_, *x*_*i*+*α*_) which is in the cycle. However, this implies *x*_*i*+*α*_ also has a cycle and *α* < ℓ because *j* > *i*; the edges from *x_i_* to *y_j_* have increasing positions. This creates a contradiction as ℓ was the *smallest* index with a cycle, so no cycles exist. Now if we have *j* < *i*, we repeat the same argument but with ℓ the largest in [0..*k* – 1] and the same contradiction arises.

The second result follows easily since clearly *M*(*i, i*), *M*(*i* + 1, *i* + 1),… are functions of sets of mutually independent random variables, and Pr(*M*(*i,i*) = 1) = (1 – *θ*).

### C.3 Proof of Lemma 2

**Supplementary Figure 2:**
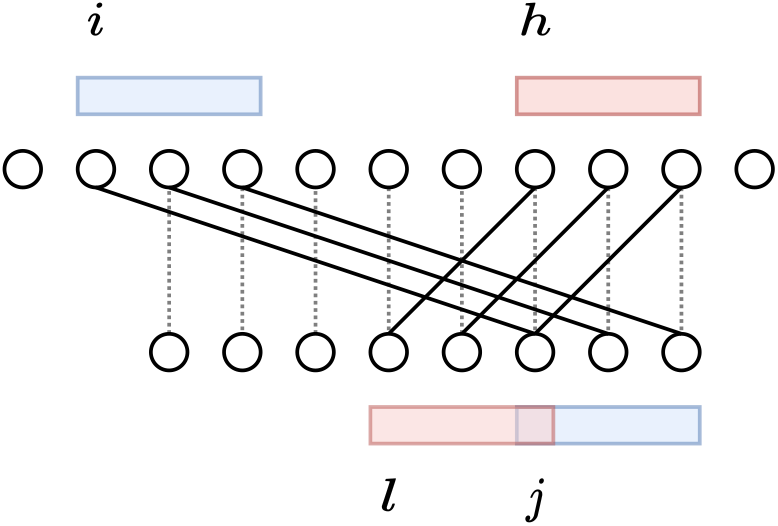
Match graph of the *M* variables induced by two anchors and pictorial representation of the setting of Lemma 2. The two anchors here generate no cycles in the match graph because |*i* – *h*| ≥ *k*, and |*i* – *l*| ≥ *k*, even though |*j* – *l*| < *k* where *k* = 3.

#### Lemma 2

*For A*(*i, j*) *and A*(*h, l*), *if both of the following conditions hold*:

1. |*i* – *h*| ≥ *k or* |*j* – *l*| ≥ k and
2. |*i* – *l*| ≥ *k or* |*j* – *h*| ≥ *k*,
*then the induced match graph on the M variables for A*(*i, j*) *and A*(*h, l*) *has no cycles*.

The intuition for the conditions of Lemma 2 is that the first condition prevents both anchors from overlapping too much, while the second condition prevents the condition where, for example, two variables *M*(1, 5),*M*(5,1) can cause a cycle to form in the match graph by *x*_1_ → *y*_5_ → *x*_5_ → *y*_5_ → *x*_1_.

*Proof.* If |*i* – *h*| ≥ *k* or |*j* – *l*| ≥ *k*, then without loss of generality, assume |*i* – *h*| ≥ *k*. Then we can find a set of {*x_i_*,…,*x*_*i*+*k*-1_} = *X_i_* and *X_h_* = {*x_h_*,…,*x*_*h*+*k*-1_} = *X_h_* disjoint. See Supplementary Figure 2 for a pictorial representation. It will turn out that if | *j* – *l*| ≥ *k*, then the xs become ys and the argument doesn’t change.

Now we claim two facts.

1. The degree of every *x* ∈ *X_i_* or *X_h_* must be at most two: each A variable induces one new edge for the xs covered by some k-mer, but these are disjoint sets of k-mers.
2. If a cycle exists, the cycle must touch some *x* ∈ *X_i_* and also a *x*′ ∈ *X_h_*: if it did not, a cycle must exist for some *x* ∈ *X_i_* or *X_h_* (the graph is bipartite, so must include a *x*) and would not use induced edges on the match graph for one of *A*(*i, j*) or *A*(*h, l*). This implies the match graph for just one of *A*(*i, j*) or *A*(*h, l*) would have a cycle, which is impossible by the proof of Corollary 1.

To complete the proof, we proceed with an argument similar to the proof of Corollary 1. We need four different cases corresponding to either |*i* – *l*| ≥ *k* or |*h* – *j*| ≥ *k* and our previous assumption of |*i* – *h*| ≥ *k* or |*j* – *l*| ≥ *k*. We prove only one case but it is easy to translate the argument to the other three.

Let’s assume |*i* – *l*| ≥ *k*. By the second fact above, if a cycle were present, we can assume it touches *x_ℓ_* ∈ *X_i_*. If *j* > *i*, let *x_ℓ_* be the leftmost *x_ℓ_* ∈ *X_i_* with a cycle, and if *j* < *i*, let *x_ℓ_* be the rightmost. Because the degree of *x_ℓ_* is at most two (by the first fact above) and thus must be exactly two, we can assume the cycle starts with the edge (*x_ℓ_, y_ℓ_*). Now, *y_ℓ_* has degree at most two since |*i* – *l*| ≥ *k*, i.e. the k-mer starting at *l* on *S*′ doesn’t overlap *y_i_*, therefore the only other edges adjacent to *y_ℓ_* from *x_α_* ∈ *X_i_*. Therefore *x_α_* is either to the left of *x_ℓ_* if *j* > *i* or to the right if *j* < *i* and must also have a cycle. However, because *x* was the leftmost or rightmost such *x*, this is impossible, so we are done.

If |*j* – *l*| ≥ *k*, we use *y*s instead of *x*s and we switch “rightmost” with “leftmost”. If we have |*j* – *h*| ≥ *k*, we switch *X_i_* or *Y_i_* with *X_h_* or *Y_h_* instead. Repeating the same argument verbatim gives the conclusion.

### C.4 Proof of Lemma 3

#### Lemma 3

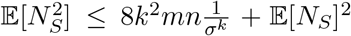. *Thus the variance Var*(*N_S_*) *can be upper bounded by* 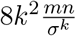. *Furthermore*, 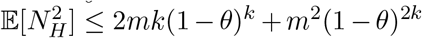 *and* 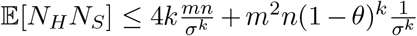.

*Proof.* We will frequently use (*m* – 1) < *m* and (*n* – 1) < *n* to simplify the bounds. With 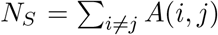, we get

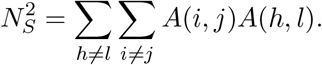

Now we try to bound 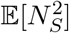, where we already know 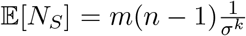. Let *B_k_*(*i, j*) = {(*h, l*) : |*h* – *i*| < *k* and |*l* – *j*| < *k*}. We can separate the sum into three different parts,

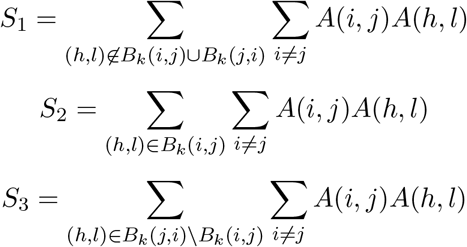

and 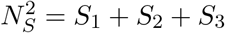. Let us bound the expectation of each sum separately. Firstly, we get

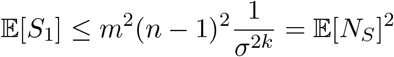

because *A*(*i, j*)*A*(*h, l*) are independent when (*h, l*) is not in the set *B_k_*(*i, j*) ∪ *B*(*j, i*) by Corollary 2, and there are at most *m*^2^(*n* – 1)^2^ possible *A*(*h, l*)*A*(*i, j*) tuples.

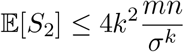

because |*B_k_*(*i,j*)| ≤ 4*k*^2^ and 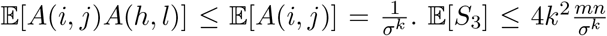 follows similarly. Thus,

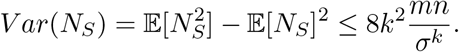

The variance of *N_H_* was calculated in (Blanca et al., 2022) for the variable *N_mut_* = *n* – *N_H_*, but we can use a simpler approximate bound. 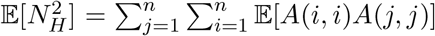; notice that if |*i* – *j*| < *k* then there is obviously dependence between *A*(*i, i*) and *A*(*j, j*) because the anchors have overlapping bases, but otherwise the variables are independent. Thus

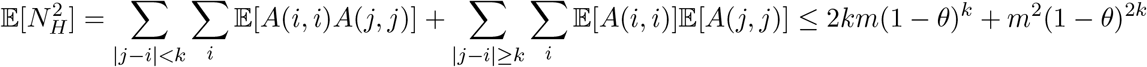

where we use the trivial bound 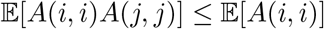 in the first bound and independence in the second.

For the bound 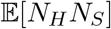, we can write this as

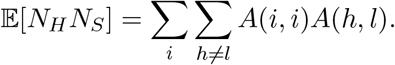

The only indices where dependence may be an issue is when either |*h* – *i*| < *k* or |*l* – *i*| < *k*. Thus for each pair *h, l*, there are at most 4*k* choices for *i* which may not be independent. We can use the same idea as the previous bound to show that

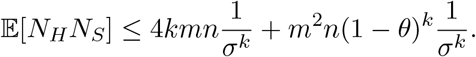

## D Missing proofs from “Proof of non-sketched main result”

### D.1 Proof of Theorem 6

#### Theorem 6

*Assume m* = *Ω*(*n*^2*Cα*+*ϵ*^) *for some ϵ* > 0, *and* 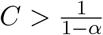. *Letting N be the total number of k-mer anchors*, 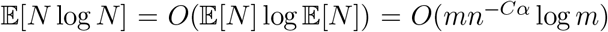. *It follows that the runtime of chaining is O*(*mn*^-*Cα*^ log *m*).

*Proof.* For all *x* > 0, we have the inequality ln(*x*/*u*) ≤ *x/u* – 1 for any *u* > 0. Substituting this into *N* ln *N* we have

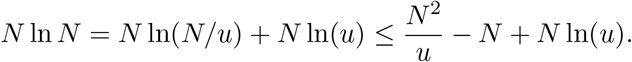

Now let 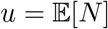 and we get 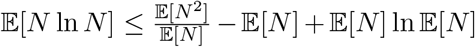. Thus if we are able to show 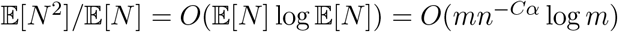 then we are done. Since 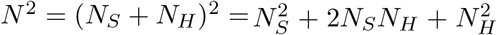, we can just use the moment bounds in Lemma 3. Plugging in the bounds yields

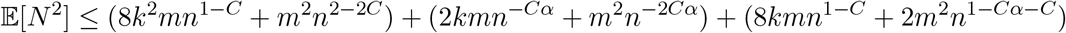

To simplify these terms, notice that *n^-Cα^* ≫ *n*^1-*C*^ and *mn^-Cα^* ≫ *k*^2^ where ≫ means asymptotically dominates. This is because –*Cα* > 1 – *C* follows from our assumption 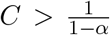. The second term is because 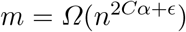 so *mn^-Cα^* ≫ *n^ϵ^* ≫ *k*^2^ = *O*(log^2^(*n*)). This gives

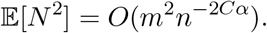

Remembering that 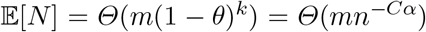, we get that 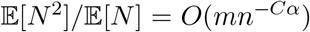 as desired, so we are done.

### D.2 Proof of Lemma 6

#### Supplementary Lemma 3.

*For any interval consisting of ℓ k-mers on S*′, *the probability that all ℓ homologous anchors are* 0 *is upper bounded by*

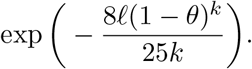

*Proof.* The letters on an interval on *S, S*′ are distributed identically as a length ℓ + *k* – 1 version of *S, S*′. We can then use Lemma 5 for *t* = *m*(1 – *θ*)^*k*^.

#### Lemma 6 (F2)

*With probability* ≥ 1 – 1/*n, no homologous gap has size greater than*

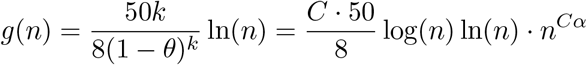

*plus a small C* log *n term we will ignore because it is small asymptotically.*

*Proof.* Using the above supplementary lemma, let 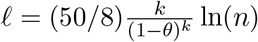. Then the probability that there are no homologous anchors in a segment of ℓ k-mers is 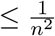.

Let *HG*_1_,…, *HG*_*m*-ℓ+1_ be indicator random variables where *HG_i_* = 1 if the next ℓ k-mers from position *i* have no homologous anchors and 0 otherwise. It follows that 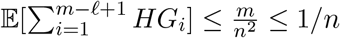. Using Markov’s inequality, we see that 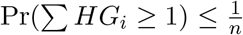. Thus with probability ≥ 1 – 1/*n*, no homologous gap is larger than 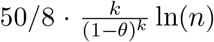 k-mers as desired. ℓ k-mers corresponds to a homologous gap of size ℓ – *k* + 1, so need to add *k* – 1 to get an upper bound on the homologous gap size, but we will ignore this in the analysis because it is small asymptotically.

### D.3 Proof of Lemma 7

#### Lemma 7 (F1 + F2)

*Take any* 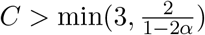 *and let* 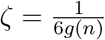 *where* 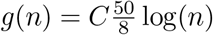 ln(*n*)*n^Cα^*. *Assume m* = *Ω*(*n*^2*Cα*+*ϵ*^) *for some ϵ* > 0. *Then for large enough n, there are no breaks of length* ≥ *m*^1/2^ *with probability greater than* 1 – 2/*n in an optimal chain*.

If we assume *C* > 3, then Lemma 4 shows that with probability ≥ 1 – 1/*n* and large enough *n*, no spurious anchors exist at all. Of course, no breaks can occur so we are already done in this case. The rest of the section is for tackling the case *C* ≤ 3. We will prove a series of supplementary lemmas, and then prove Lemma 7.

We will now assume the hypotheses of Lemma 7 for the rest of this section. That is, 3 ≥ *C* > 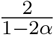, *m* = *Ω*(*n*^2*Cα*+*ϵ*^) for some *ϵ* > 0, and define 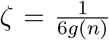 as in the statement of Lemma 7. We will not be too careful with small additive constants of order *O*(log *n*) due to indexing offsets from now on. For example, a length *m*^1/2^ interval of bases contains technically *m*^1/2^ – *k* + 1 k-mers, but since we work with asymptotics we’ll treat this as ~ *m*^1/2^.

#### Supplementary Lemma 4 (F1 + F2).

*With probability* ≥ 1 – 2/*n, given an optimal chain* ((*i*_1_, *j*_1_),…, (*i_u_, j*_u_)) *at least one of the anchors is a homologous anchor for large enough n*.

*Proof.* We will show that no chain of only spurious anchors can be optimal, implying that any optimal chain must have at least one homologous anchor. By F2, there are no homologous gaps of length ≥ *g*(*n*) where *g*(*n*) = *Θ*(*n^Cα^* log^2^(*n*)), so we can lower bound the number of homologous anchors *N_H_* under the space F2 by

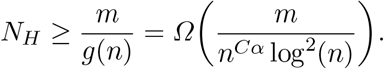

Let score(*N_H_*) be the score of the chain with only homologous anchors. Then

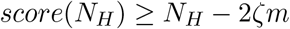

because the only homologous chain has a linear cost of at most 2*ζm*. Now consider a chain without homologous anchors, i.e. only spurious anchors. Such a chain has maximum score

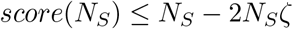

because there are at most *N_S_* such anchors, and the *N_S_* in linear cost is assuming the smallest possible linear cost where there are no gaps between the anchors, i.e. (*i_u_* – *i*_1_) + (*j_u_* – *j*_1_ ≥ 2*N_S_* if *u* = *N_S_*. Using condition F1, 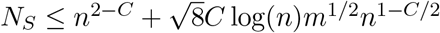. Since 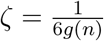,

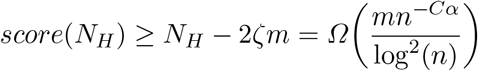

holds and also

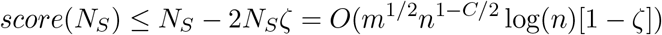

holds. We therefore we want the following asymptotic inequality to hold:

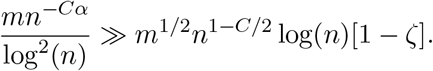

This holds when 1 – *C*/2 + *Cα* < 0 which is equivalent to our assumed condition 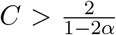, and thus the chain of only homologous anchors has asymptotically greater score than any chain with only spurious anchors. Therefore any optimal chain with high probability must contain at least one homologous anchor for large enough *n*.

#### Supplementary Lemma 5 (F1 + F2).

*Suppose a break is flanked by two homologous anchors in an optimal chain. That is, homologous anchors exist on both sides of the break in the chain. Then with probability* ≥ 1 – 2/*n and large enough n, this break has size* < *m*^1/2^.

*Proof.* Suppose a break of length ≥ *m*^1/2^ is flanked by two homologous anchors. Then for the given chain ((*i*_1_, *j*_1_),…, (*i_u_, j_u_*)), the break occurs somewhere in the middle, say ((*i_r_, j_r_*),…, (*i_t_, j_t_*)) where *r* > 1, *t* < *u*. Let us construct a new chain by removing all spurious anchors ((*i_r_, j_r_*),…, (*i_t_, j_t_*)) within our chain that lie in this break, and then adding all homologous anchors that are present within the break to the chain. See Supplementary Figure 3.

This does not change the gap cost part of the chaining score which is still *ζ*[(*i_u_*–*i*_1_) + (*j_u_*–*j*_1_)], so if there are more homologous anchors after the switch, then this switch is more optimal. Assuming condition F2, the number of homologous anchors contained within a break is lower bounded by *m*^1/2^/*g*(*n*). Assuming the condition F1, the number of spurious anchors, and therefore the size of the break, is upper bounded by *N_S_* = *Θ*(log(*n*)*m*^1/2^*n*^1-*C*/2^). Notice that

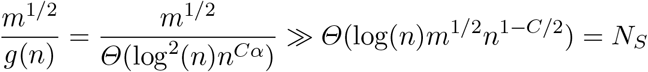

again exactly when 1 – *C*/2 + *Cα* < 0 which is equivalent to 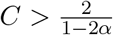. Thus under the event space F1 ⋂ F2, switching to the homologous k-mers always improves our optimal chain for large enough *n*.

#### Supplementary Lemma 6 (F1 + F2).

*Suppose a break is flanked by a single homologous anchor in an optimal chain. Then with probability* ≥ 1 – 2/*n and large enough n, this break has size* < *m*^1/2^.

*Proof.* It follows easily from the definition of a break that a break flanked by one homologous anchor must occur at the start or end of the chain ((*i*_1_, *j*_1_,…, (*i_u_, j_u_*)). Let us assume that such a break occurs at the end of the chain at anchors ((*i_r_, j_r_*),…, (*i_u_, j_u_*)); the same argument works for the break occurring at the beginning of the chain.

We use a similar argument as above, but this time we must take into account the linear gap cost. Furthermore, instead of switching to the relative homologous anchors, we just remove the break from the chain and check if that improves the score. See Supplementary Figure 4.

**Supplementary Figure 3:**
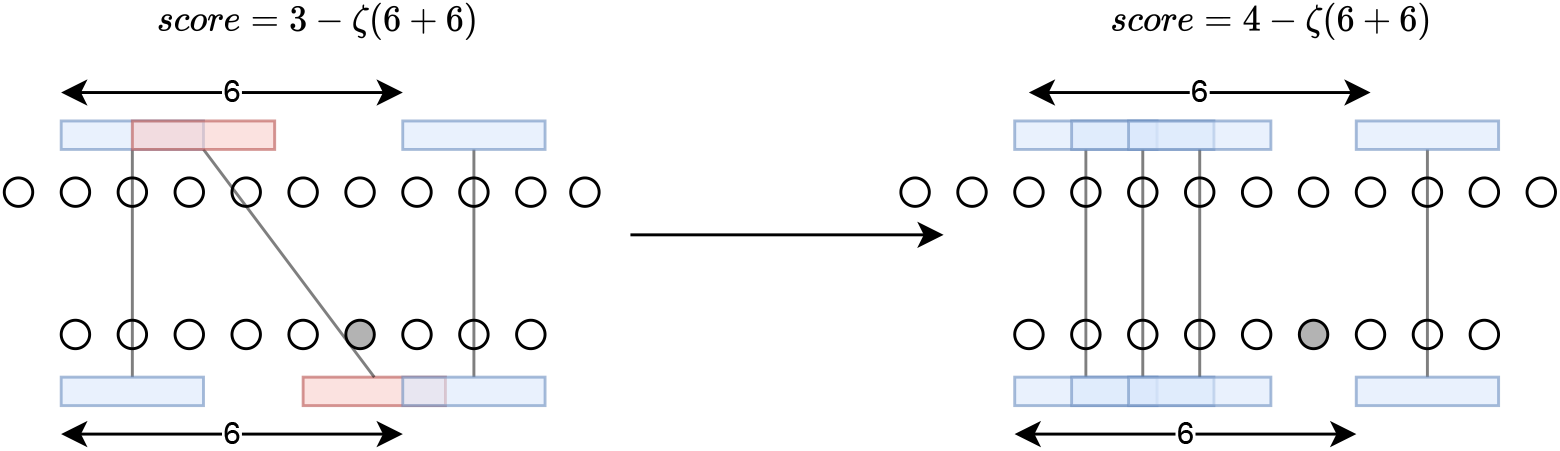
Circles are bases, grey circles are mutated bases, and boxes are k-mers. The red spurious anchor indicates a break. We can always remove a break flanked by two homologous anchors and then add in homologous anchors that may be present within the break. This idea is used in the proof of Supplementary Lemma 5 to show that for large enough breaks, such a procedure always improves the chaining score.

**Supplementary Figure 4:**
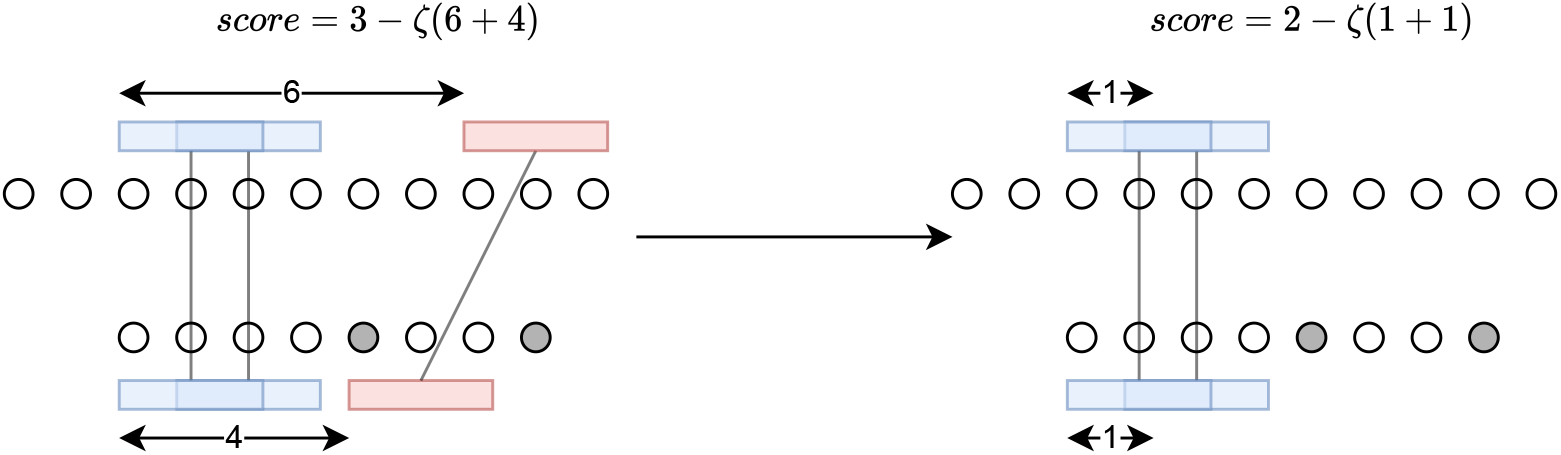
Removing a break (the red anchor of k-mers) at the end may increase the chaining score if the break is long enough. We use this idea in the proof of Supplementary Lemma 6 to show that large breaks can not appear near the ends of chains.

The score of the old chain is *A* – *ζ*[(*i_u_* – *i*_*r*-1_) + (*j_u_* – *j*_*r*-1_)] + *w* where *A* is the score considering subchain of the anchors prior to *r*, and *w* is the number of anchors in the break. This is upper bounded by *A*–*ζ*(*B* – *k*)+*N_S_* where *N_S_* = *Θ*(log(*n*)*m*^1/2^*n*^1-*C*/2^) as before and *B* is the length of the break. This holds because the length of the break subtracted by *k* is *B* – *k* = max(*i_u_, j_u_*) –min(*i_r_, j_r_*) which is less than [(*i_u_* – *i*_*r*-1_) + (*j_u_* – *j*_*r*-1_)]. Removing the break entirely is more effective when *ζB* – *k* > *N_S_*, so when

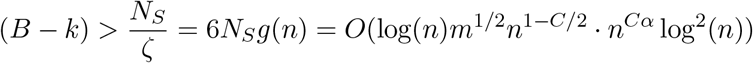

we can get a better chain by simply removing the break. By the previous discussion, 1 – *C*/2 + *Cα* < 0, so we just let *B* = *m*^1/2^ and this completes the proof.

#### Proof (Lemma 7).

Breaks are flanked by either one, two, or no homologous anchors. The last case can only occur if the entire chain is a break. By Supplementary Lemma 4, we always have at least one homologous anchor in any optimal chain (under F1 ⋂ F2), so any break is flanked by at least one homologous anchor. Therefore by the previous supplementary lemmas, no breaks of size *m*^1/2^ occur with probability ≥ 1 – 2/*n* after using the conditions F1, F2 with the appropriate assumptions.

### D.4 Proof of Corollary 3

#### Supplementary Lemma 7.

*Under the assumptions of Lemma 7, given any optimal chain* (*i*_1_, *j*_1_),…, (*i_u_, j_u_*), 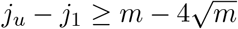 *with probability* ≥ 1 – 2/*n for large enough n*.

*Remark 1.* The value 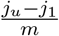 is called the aligned fraction and is clearly an upper bound on recoverability. The above Supplementary Lemma shows that the expected value of the aligned fraction is 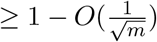.

*Proof.* Lemma 7 shows the max break size is *m*^1/2^; we claim that 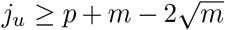, i.e. *j_u_* is less than 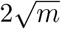 away from the end of *S*′. Suppose otherwise. The argument proceeds in two cases. The idea is to essentially show that adding on all of the homologous anchors near the end of *S*′ always increases the score (under the event space F1 ⋂ F2).

If (*i_u_, j_u_*) is homologous, then *m*^1/2^ ≫ *g*(*n*): *g*(*n*) = *Θ*(*n^Cα^* log^2^(*n*)) is the maximum distance between homologous k-mers and *m* = *Ω*(*n*^2*Cα*+*ϵ*^). Thus for large enough *n*, we can find another homologous k-mer near the edge of *S*′. Adding this homologous k-mer to the chain changes the score by at least –2*ζg*(*n*) + 1 where the –2*ζg*(*n*) is the linear cost. Since 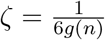, this is positive, so the old chain was not optimal.

**Supplementary Figure 5:**
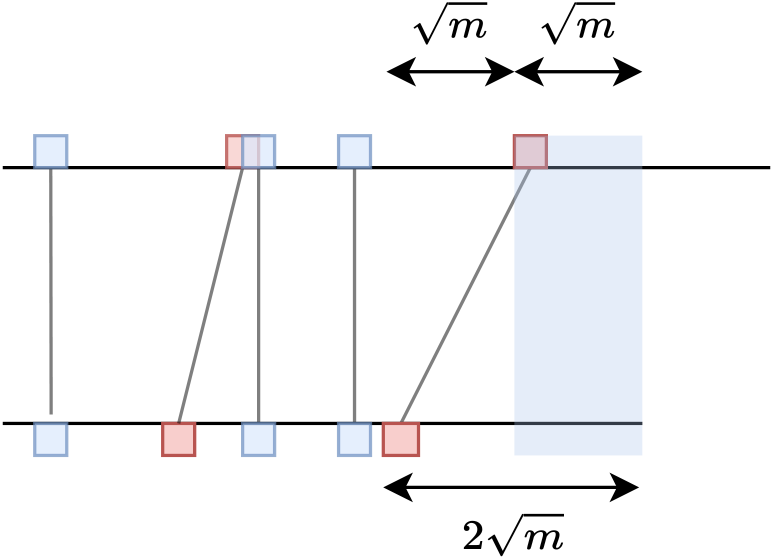
Given an optimal chain (shown with k-mer anchors in red and blue), if the last k-mer on *S*′ is 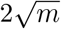 distance away from the end, because the break size is 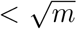, there will remain at least 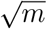 bases left (shaded in blue) near the end of *S*′, *S*. We use this geometry in the proof of Supplementary Lemma 7 and argue that adding in every possible homologous k-mer in the shaded region gives a more optimal score.

For the non-homologous case, refer to Supplementary Figure 5 for the geometry. Now if (*i_u_, j_u_*) is not homologous, then 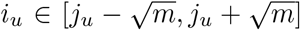 because any spurious anchor is contained in a break, which has size 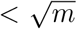. Assume *j_u_* = *p* + *m* – *L* where 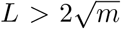 so *j_u_* is *L* away from the end of *S*′.

We claim that adding in all homologous anchors in the interval [max(*i_u_, j_u_*) + 1..*p* + *m*] gives a more optimal chain. Indeed, since 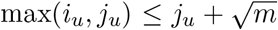, 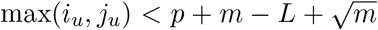, and thus there are at least 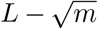 homologous positions to the right of max(*i_u_, j_u_*). Thus there are at least 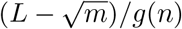 additional homologous anchors.

The additional linear cost from adding on these homologous anchors near the end is at most *ζ*[(*p* + *m*) – (*p* + *m* – *L*)] = *ζL* on the side of *S*′ by lengthening the chain from *p* + *m* – *L* to *p* + *m*, and at most 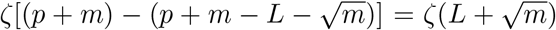 on the side of *S* for the same reason. Summing these two terms gives 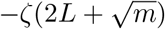 as the maximal linear cost penalty of adding these new anchors. However,

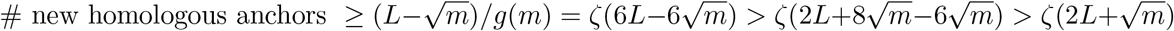

after using the inequality 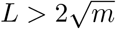. Therefore adding in these homologous k-mers makes the score bigger, contradicting the optimality of our chain. Thus 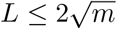, and the distance from *j_u_* to the end of *S*′ is at most 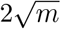.

The argument works with directions flipped for *j*_1_, so 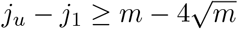 as desired.

#### Corollary 3

*Under the same assumptions as in Lemma 7, the expected recoverability of any optimal chain is* 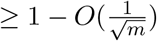 *for large enough n*.

*Proof.* Let 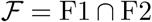, so all breaks have length 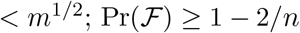. Letting 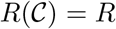 be the recoverability as before and letting |*S*′| = *m* + *k* – 1 ~ *m* because our final result uses big O notation anyways, we get from Lemma 1 relating recoverability to breaks that

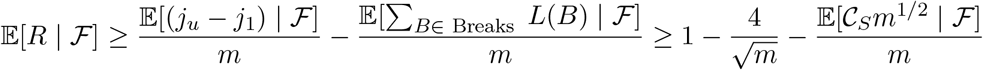

where we define 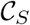 to be the number of spurious anchors in an optimal chain. The number of breaks is upper bounded by 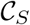, and we’ve used Lemma 7 and Supplementary Lemma 7 to bound *L*(*B*) by *m*^1/2^ and *j_u_* – *j*_1_ by 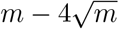 respectively. It is clear that 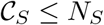 so

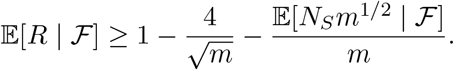

Since 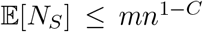, we have that 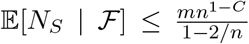 by the law of total expectation and non-negativity of *N_S_* Using this, we get that

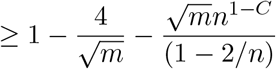

and 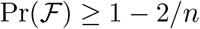, so

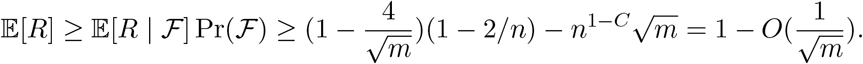

*Remark 2.* In the proof of the intermediate lemmas associated with the proof of Corollary 3, we assumed that *S*′ was not too close to the edges of *S*, i.e. quantities such as 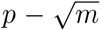 were nonnegative. It’s not hard to see that if *p* were close to the edges of S, the breaks would actually be smaller because by definition breaks can not go past the ends of *S*. One could in fact show that our upper bound still work after accounting for these edge cases, but for simplicity, we omit these edge cases from our proof.

### D.5 Proof of Lemma 8

We prove Lemma 8 by proving a series of lemmas.

#### Supplementary Lemma 8.

*For a given instance of S,S′, let* 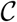 *be any optimal chain. Let* 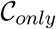 *be the chain consisting of only homologous anchors, in S,S*′. *Defining* 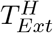 *to be the runtime of extension of* 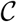 *over only the homologous gaps in* 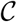 *and* 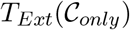 *as the runtime of extension over the homologous gaps of* 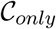, *we have that*

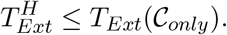

*Proof.* Let (*i_ℓ_, i_ℓ_*) and (*i*_ℓ+1_, *i*_ℓ+1_) be two consecutive homologous anchors in an optimal chain 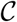. We can guarantee that no *A*(*i_ℓ_* + 1, *i_ℓ_* + 1),…, *A*(*i*_ℓ+1_ – 1, *i*_ℓ+1_ – 1) random variables are equal 1, otherwise adding such an anchor would improve an optimal chain (it does not incur a linear gap cost penalty due to being flanked by two anchors). Thus (*i_ℓ_, i_ℓ_*) and (*i*_ℓ+1_, *i*_ℓ+1_) are also consecutive anchors in 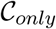, and the homologous gap corresponding to those anchors is also present in 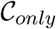. Therefore, 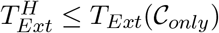 since all homologous gaps in 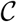 are also in 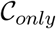.

#### Definition 6.

*Define* 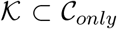 *to be the subchain of homologous anchors for which the starting positions of the k-mers on S are in* 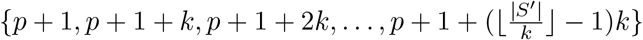.

That is, the subchain 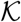 consists of only homologous anchors restricted to k-mers that are spaced *k* bases apart starting from the first index. Since 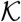 is sparser than 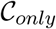, it should take longer to extend through 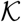 than 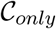. We formalize this below.

**Supplementary Figure 6:**
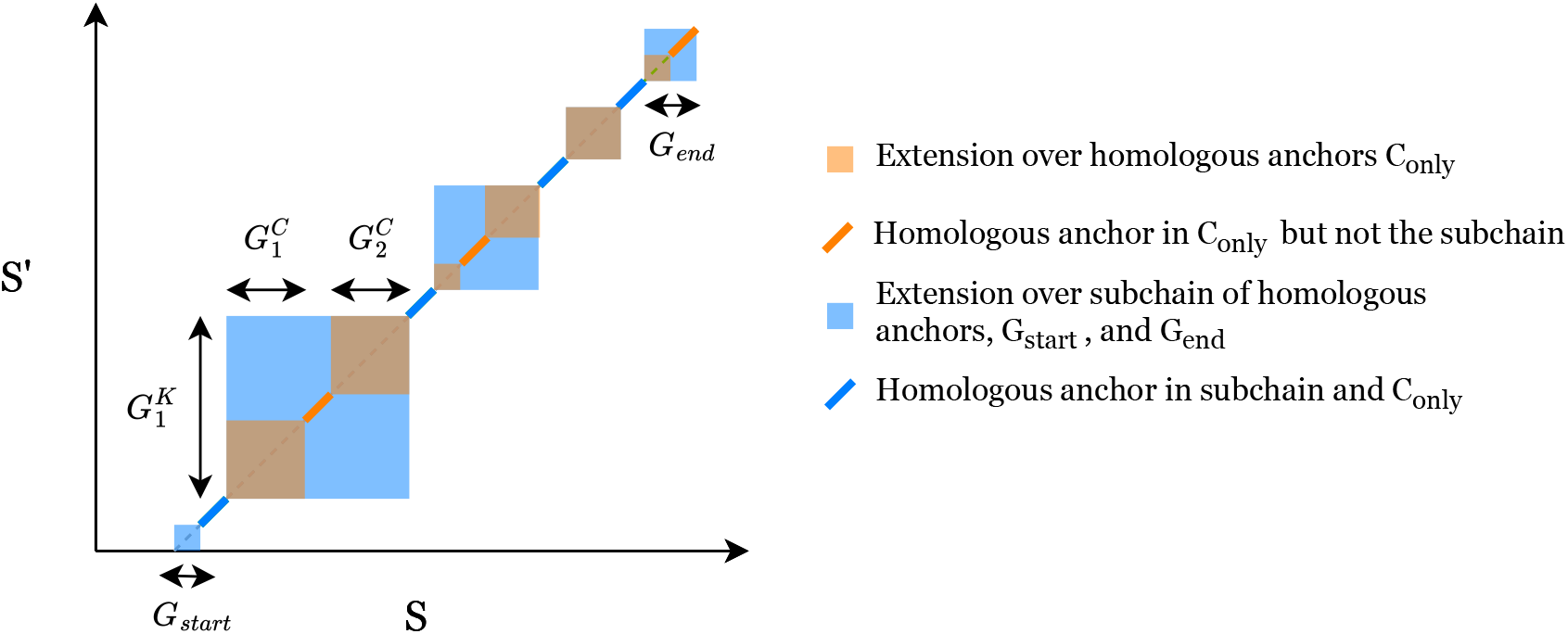
A graphical proof of Supplementary Lemma 9. Given a subchain of homologous anchors, the extension time is longer because the square of the larger gaps in the subchain contains the square of the smaller gaps over all homologous anchors. The last gap on the right is not accounted for in the subchain, but adding 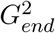 and 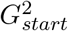 fixes this. Note: the second last orange square on the right is a gap for both chains, i.e. the square is both orange and blue.

#### Supplementary Lemma 9.

*Let G_start_ be the distance from the first anchor of* 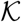 *to the start of S*′ *and similarly for G_end_ and the last anchor of* 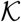 *to the end of S*′. *Then*

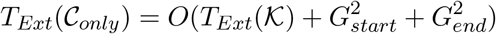

*Proof.* See Supplementary Figure 6 for a visualization of the proof. Let 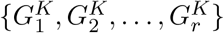 be the homologous gap sizes in 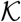, where we think of 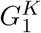 as the gap size (possibly 0) between the first two anchors, and so forth. Similarly, let 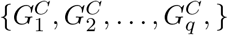 be the homologous gap sizes in 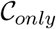. The extension runtime is

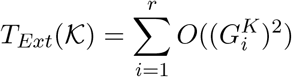

and similarly for 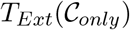. Any two consecutive anchors in 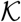 give rise to a gap 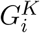. These two anchors also exist on 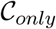, but there may be intermediate anchors, so this gives rise to multiple gaps 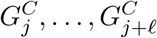 between these two anchors on 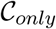. Now 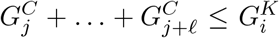 because the sum of all intermediate gaps is at most the size of the larger gap, so

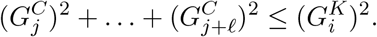

Let 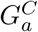 be the leftmost gap on 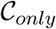 to the right of the first anchor of 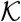, and 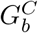 be the gap to the left of the last anchor of 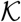. Considering every pair of consecutive anchors on 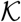, we get

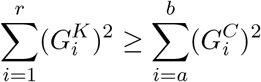

by the previous inequality. However, we’re not done yet because the leftmost gaps in *C_only_* may not be contained by any gap in 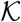. Using the definition of *G_start_* as the “gap” on 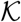 containing all of the leftmost gaps on 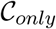 and similarly for *G_end_*, we get

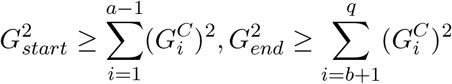

by the same arguments as above. Combining both inequalities gives the result.

By the above results, we can bound 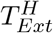 by either 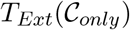 or 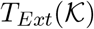. We will work with 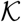 in this section, but we will actually use 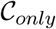 for the sketched version of the main theorem. Below we give a random variable to calculate these extension times.

#### Definition 7.

*Let Y_i_ be the random variable representing the number of uncovered bases between a homologous anchor starting at position i, if it exists, and the next homologous anchor. So Y_i_* = ℓ ≥ 1 *if the bases* [*i..i* + *k* – 1] *are unmutated, the bases at* [*i*+ℓ + *k*..*i*+ℓ + 2*k* – 1] *are unmutated, and there are no homologous anchors covering any bases in* [*i* + *k..i* +ℓ + *k* – 1]. *Otherwise, Y_i_* = 0. *Let* 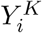 *be the same random variable except only considering homologous anchors with starting positions in* 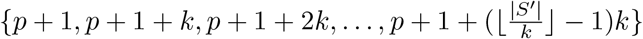.

Under our original definition, the extension time over 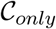 would be 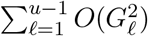 where *G_ℓ_* is the size of the gap, but *u* is a random variable. It’s clear that 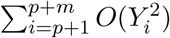 is the runtime of extension over 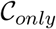, but it will be easier to handle for our proof now that the upper index is not a random variable. Similarly, 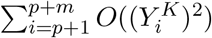 is the runtime of extension over 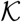.

We will now work with the chain 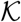 restricted to equally spaced k-mers and their unmutated homologous anchors. We upper bound this extension time in expectation, which will upper bound 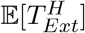 as well.

#### Lemma 8

*Let* 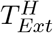 *be the time of extension over only the homologous gaps of an optimal chain*. 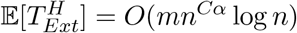.

*Proof.* By Supplementary Lemmas 9 and 8, to bound 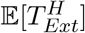 we can bound 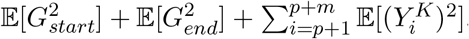. The random variable *G_start_* = ℓ · *k* if for the first ℓ k-mers, *A*(1,1) = 0, *A*(1 + *k*, 1 + *k*) = 0,…, *A*(1 + (ℓ – 1)*k*, 1 + (ℓ – 1)*k*) = 0 but *A*(1 + ℓ*k*, 1 + ℓ*k*) = 1. In other words, the first ℓ k-mers that are spaced *k* bases apart are mutated, but the ℓ + 1th such k-mer is not mutated. Thus

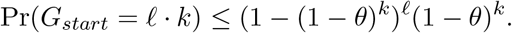

Importantly, we used the fact that k-mers spaced *k* distance apart are independent of each other. The ≤ comes from the fact that *G_start_* can not be larger than |*S*′|. It follows that

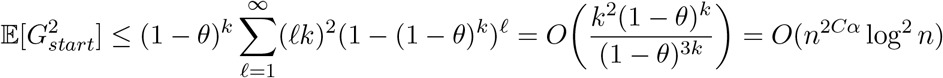

by the formula

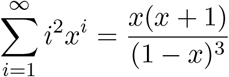

which follows from geometric series manipulations. Notice that *O*(*n*^2*Cα*^ log^2^ *n*) = *O*(*m*) because *m* = *Ω*(*n*^2*Cα*+*ϵ*^). It’s clear that the same argument holds for 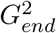, so both terms are *O*(*m*).

Now for 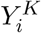, we have that 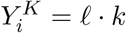 where ℓ > 0 only if *i* ∈ {*p* + 1, *p* + 1 + *k, p* + 2 + *k*,…}, the k-mer at *i* is unmutated, the k-mers that are *k*, 2*k*, 3*k*,…,ℓ*k* bases ahead from *i* are mutated, and the k-mer (ℓ + 1) · *k* bases ahead of i is unmutated. Thus

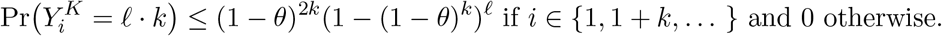

We can compute the expectation over all 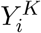 and only pick out the non-zero random variables where *i* ∈ {*p* + 1, *p* + 1 + *k*,…}. There are at most *m/k* such random variables, so we get

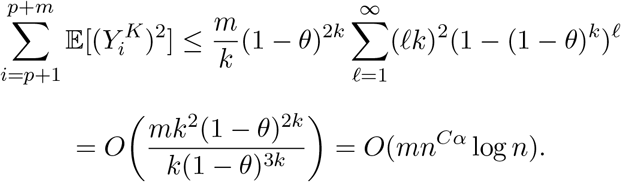

The expectations in all the terms are *O*(*mn^Cα^* log *n*), so this finishes the proof.

### D.6 Proof of Lemma 9

#### Supplementary Lemma 10 (F1 + F2).

*Under the same assumptions of Lemma* 7 *all non-homologous gaps in an optimal chain have size* 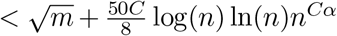 *on both S and S*′ *with probability* ≥ 1 – 2/*n and large enough n*.

*Proof.* Let us be in the space 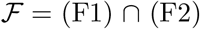, which holds with probability ≥ 1 – 2/*n*. There are two types of non-homologous gaps; non-homologous gaps that are flanked by two spurious anchors and gaps that are flanked by only one.

If a non-homologous gap is flanked by two spurious anchors, then it is part of a break. The gap size must be smaller than the break, which has length less than 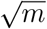.

Suppose a non-homologous gap is flanked by one spurious anchor *A*(*i, j*) and one homologous anchor *A*(*h, h*). Suppose WLOG that |*i* – *h*| > |*j* – *h*|. We know with high probability that |*j* – *h*| < *g*(*n*) = 50/8C log(*n*) ln(*n*)*n^Cα^* as otherwise there will be a homologous anchor between *A*(*i, j*) and *A*(*h, h*) (by property F2/Lemma 6) and the chain is not optimal as we could just insert the additional homologous anchor between *j* and *h*. Furthermore, 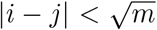 as |*i* – *j*| is less than the break size. We can see then that 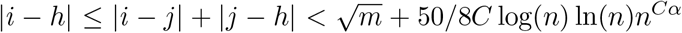 as desired.

#### Lemma 9

*Let* 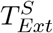 *be the runtime of extension through only the non-homologous gaps of an optimal chain. Under the same assumptions as in Lemma 7*, 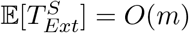.

*Proof.* Defining *γ* as the maximum non-homologous gap size as in Supplementary Lemma 10 conditional on 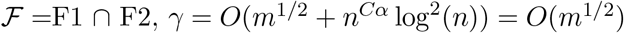 when *Ω* = (*n*^2*Cα+ε*^). We bound the expected value of the non-homologous gaps as follows.

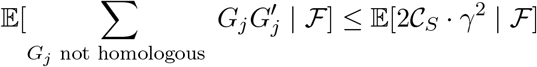

The inequality uses *γ* > *G_j_* and that the number of non-homologous gaps is at most 2 times 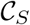, which we define to be the number of spurious anchors in the chain (each anchor gives rise to at most two unique gaps). 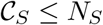 follows trivially, so

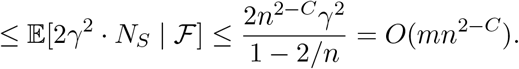

The first inequality follows from 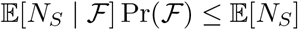 and non-negativity of *N_S_*, as well as 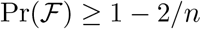. Finally, under 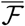, the worst case extension through non-homologous gaps is just *O*(*nm*) as in section “Extension and chaining runtimes”. Since 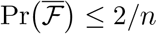, the expected runtime is *O*(*nm*/*n* + *mn*^2-*C*^) = *O*(*m*) as desired.

## E Missing proofs from “Sketching and local k-mer selection”

### E.1 Proof of Lemma 11

#### Lemma 11

*The variance* 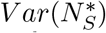 *can be upper bounded by* 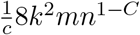. *Furthermore*, 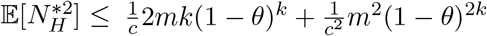 and 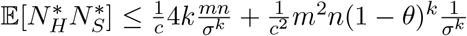.

*Proof.* The proof follows almost exactly the same as Lemma 3. We only do the variance bound as an example, and the other moment bounds follow exactly the same way.

We upper bound the three sums *S*_1_, *S*_2_, *S*_3_ in the proof of Lemma 3 but now with the sketched versions 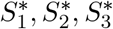. The bounds 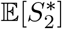 and 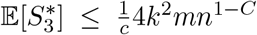 hold by a restatement of the argument using 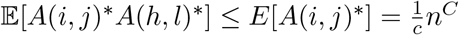 and the set |*B_k_*(*i,j*)| ≤ 4*k*^2^.

The bound for 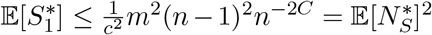 also holds as well; we just have to calculate 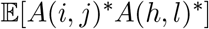 when *A*(*i,j*) and *A*(*h,l*) are independent. This is

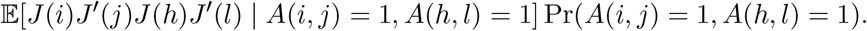

By independence of *A*(*i, j*), *A*(*h,l*) in 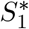, the second term is *n*^-2*C*^. Now notice that under the conditions of *S*_1_, either |*i* ≥ *h*| > *k* or |*j* – *l*| ≥ *k* meaning that two of the k-mers along either *S* or *S*′ are independent. WLOG we can assume it is *i* and h; thus *J*(*i*) and *J*(*h*) are independent. The *A*(*i,j*) = 1 condition implies *J*(*i*) = *J*′(*j*) and similarly for *h,l*, so

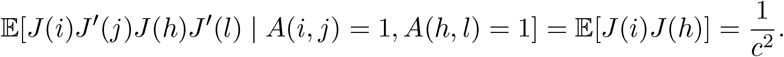

### E.2 Proof of Theorem 8

#### Theorem 8

*Under the same assumptions as in Theorem 6, letting N* be the total number of sketched k-mer anchors, the expected chaining time is* 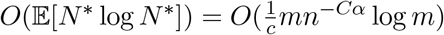.

*Proof.* The argument follows in the same way as in the proof of Theorem 6, only using the sketched moment bounds in Lemma 11. We still get that 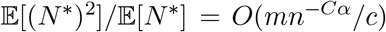 and the result follows.

### E.3 Sketched concentration bounds and gap sizes

#### Supplementary Lemma 11.

*Let τ* = (*c* + 1)/2 *and* 1 > *β* ≥ 0. *Given ℓ* + *τ* – 1 *k-mers indexed by the interval* [*a..b*] *on a random string S*,

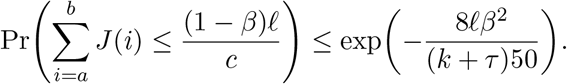

*Remark 3.* The variables *J*(*i*) are k-dependent so we could use Theorem 5 for this sum. This however leads to a bound like exp(–*O*(*m*/(*kc*))) instead of our bound which is exp(–*O*(*m*/(*k* + *c*))) in the above lemma. While the former bound still leads to the same asymptotic behavior for Supplementary Lemma 13, which we are ultimately trying to prove, we believe it is enlightening to show how the dependence structure of the *J*(*i*)s can be used to obtain a better bound.

*Proof.* Define *J*(*i,i* + *τ*) = *J*(*i*) + ⋯ + *J*(*i* + *τ* – 1). Then

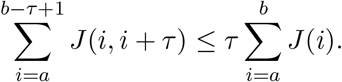

Let min(*J*(*i, i* + *τ*), 1) = *I*(*i, i* + *τ*) and clearly 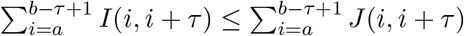.

We will show that 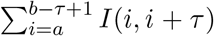 is large, implying that 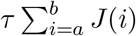 is also large. Notice that *I*(*i, i* + *τ*) is the random variable which is 1 when some k-mer in [*i..i* + *τ*) is an open syncmer. By Theorem 7, only one of the *J* variables in *J*(*i*),…, *J*(*i* + *τ* – 1) can be equal to one. Using 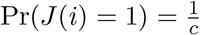, we get that

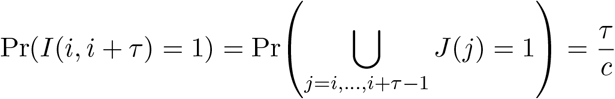

by disjointness. *I*(*i, i* + *τ*) is *k* + *τ* – 1-dependent because it examines the *k* + *τ* – 1 bases starting from position *i*, so using Theorem 5 we get that

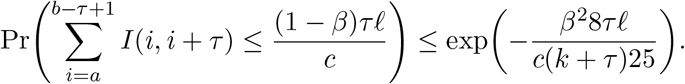

From our inequality 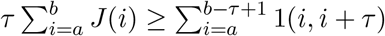 and using *τ* = (*c* + 1)/2 we obtain

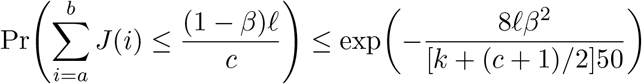

#### Supplementary Lemma 12.

*For any interval* [*a..b*] *containing* 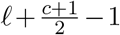 *k-mers and* 1 > *β* ≥ 0, *the probability that all A*\*(*a, a*),…, *A*\*(*b, b*) *are* 0 *is upper bounded by*

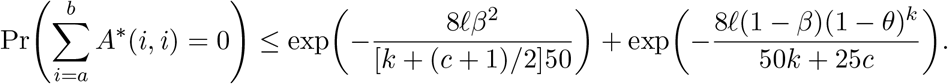

*Proof.* Defining 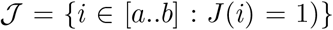, we have 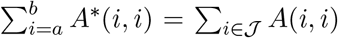 where we just sum only over syncmer anchors instead of all k-mer anchors in the range.

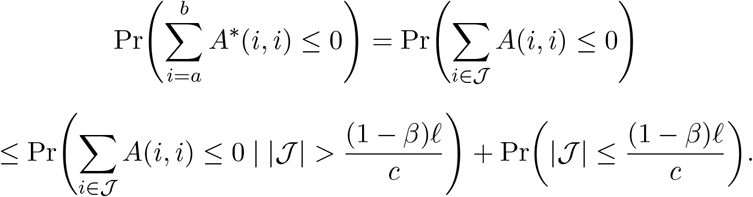

Now by Theorem 7, the distance between consecutive positions in 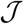 is at least (*c*+1)/2 (remembering that we assume *c* is odd for simplicity), which means that given the ith open syncmer, the 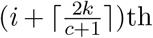 open syncmer is more than *k* bases apart from the ith open syncmer and hence independent from each other. Thus the *A*(*i, i*)s in 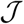 are now 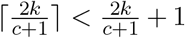 dependent. Conditioning on 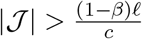, we can bound the first term using Theorem 5 to get

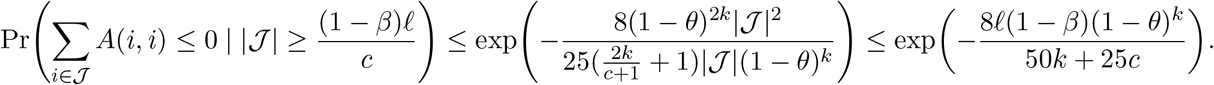

Where we substituted in 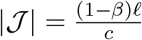 in the above equation and used 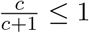 to remove the term. To bound the second term, we just use Supplementary Lemma 11. This furnishes the final result

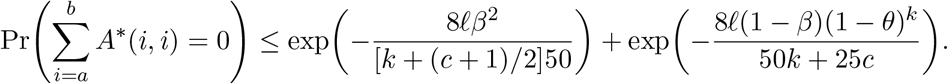

#### Supplementary Lemma 13.

*No homologous gap has size greater than*

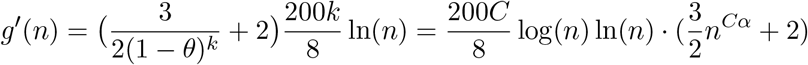

*with probability* ≥ 1 – 2/*n after ignoring an additive* (*c* + 1)/2 – 1 + *C* log *n term.*

*Proof.* We use Supplementary Lemma 12 after plugging in the value of ℓ = *g*′(*n*) and letting *β* = 1/2. Algebraic manipulations show that the probability is less than ≤ 2/*n*^2^ using *k* = *C* log *n* and the inequality *c* < *k*. Then as in the proof of Theorem 6 we can use indicator random variables and Markov’s inequality in the same way to get the result.

### E.4 Proving Lemma 13

We use the same definitions and supplementary lemmas as in Section D.5 for *Y_i_*, 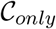 and 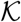, now considering the sketched versions 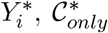, and 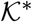. We first note that the sketched version for Supplementary Lemmas 8 and 9 hold by the same arguments. The analogous version of Lemma 8 holds as well.

#### Supplementary Lemma 14.

*Let* 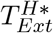 *be the time of sketched extension over only the homologous gaps of an optimal chain*. 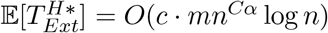.

*Proof.* Essentially the same argument as in the proof of Lemma 8 outlined in Section D.5. We end up bounding 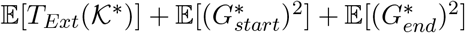; the only difference is that 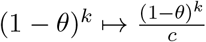 due to sketching. Ultimately, the main term becomes

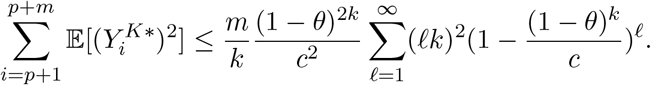

The same geometric series manipulation as before gives us that this is

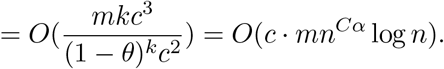

This finishes the proof.

When *c* is fixed to be a constant independent of *n*, we get the same asymptotic bound as before. This bound suggests that sketching makes extension slower. However, we can actually do better than this when *c* grows with *n*. This time, we proceed in a different manner, instead directly using 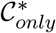 to bound the extension time. We let 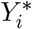 be the sketched version of *Y_i_* as in Definition 7, which is a random variable measuring gap sizes between anchors.

#### Supplementary Lemma 15.

*For* ℓ ≥ 1,

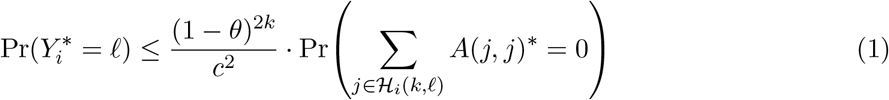

*where* 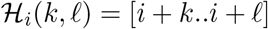 *if* ℓ ≥ *k, and is empty otherwise. Thus* 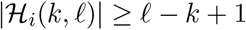.

*Proof*. 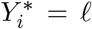 is equivalent to the k-mer covering [*i..i* + *k* – 1] being unmutated (i.e. no bases on the k-mer are mutated) and selected as an open syncmer, all of the k-mers in between these two flanking k-mers not being present, and the k-mer covering [*i* + ℓ + *k..i* + ℓ + 2*k* – 1] being unmutated and selected as an open syncmer. Calling these events *H*_1_, *H*_2_, and *H*_3_ respectively, Pr(*Y_i_* = ℓ) = Pr(*H*_1_ ⋂ *H*_2_ ⋂ *H*_3_)

The k-mers considered in *H*_2_ that lie in between the flanking k-mers overlap the flanking k-mers in *H*_1_ and *H*_3_, so these events are not independent. To upper bound *H*_2_, let 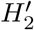 be the event that the k-mers lying *completely in the interval* [*i* + *k..i* +ℓ + *k* – 1] on *S*, and not just overlapping the interval, are mutated or not selected as an open syncmer. 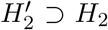 as events. This set of k-mers is exactly the k-mers starting at positions in 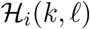 after some examination. Now notice that *H*_1_, 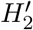, *H*_3_ are all independent as the k-mers in each event lie on non-overlapping bases, so

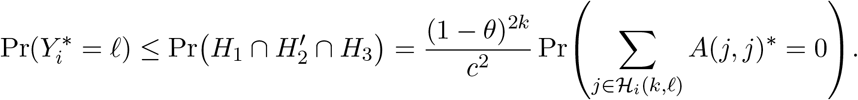

In the case that [*i* + 1 + *k..i* + 1 + 2*k* – 1] does not lie on *S*′, i.e. *i* + 1 + *k* > *p* + *m*, then Pr(*Y_i_* = ℓ) = 0 as a homologous gap of this length can not exist near the edges, so the upper bound still holds. The upper bound also holds if *i* – *k* + 1 < *p* + 1 by the same reason. This finishes the proof.

#### Lemma 13

*The expected runtime of sketched extension through the homologous gaps in an optimal chain is* 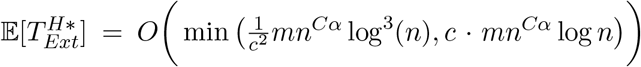. *If* 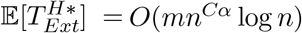.

*Proof.* By Supplementary Lemma 12 and our discussion that 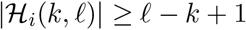,

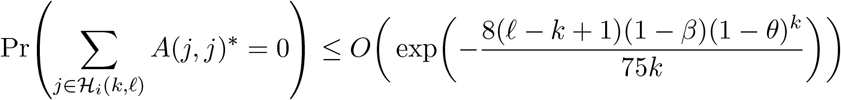

holds by noticing that the first term in Supplementary Lemma 12 is dominated by the second term and using *k* > *c*, where we also ignored the 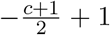 in Supplementary Lemma 12 because it is asymptotically small. We can now bound the expected time of extension over homologous gaps by 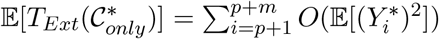. Since the probability densities of each 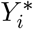 is upper bounded by Equation 1,

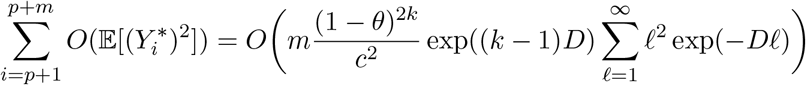

where 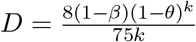 and *β* is some constant. Now we use the fact that

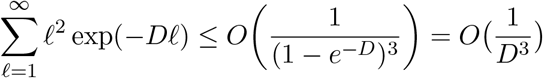

after geometric series manipulations as before, using Taylor’s theorem for 1 – *e*^-*D*^, and noticing that *D* = *o*(1). Thus our final bound is

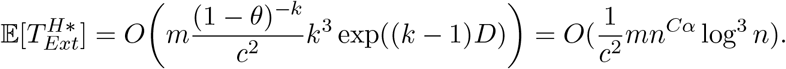

The other bound *c* · *mn^Cα^* log *n* follows from Supplementary Lemma 14, so taking the minimum over both bounds yields the result.

*Remark 4.* Intuitively, what is happening here is that open syncmers have the same key property as K; they are spaced at least 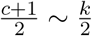 bases apart when *c* ~ *k*, and are “almost independent” like the anchors in 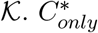 has similar properties to the non-sketched 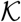 chain, showing why they give the same bounds. Interestingly, if were to bound 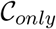 to get a result for the non-sketched version 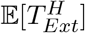 while using the same techniques as above (with the dependent Chernoff-Hoeffding bound), the result would be *O*(*mn^Cα^* log^3^ *n*), worse than bounding using 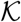 by log^2^(*n*).

*Remark 5.* If we were to use the FracMinHash method or another k-mer selection/sketching method without the distance guarantee provided by Theorem 7, Supplementary Lemma 12 would not hold, and therefore the above argument would fail. Supplementary Lemma 14 would however still hold.

### E.5 Re-proving sketched bounds

#### Supplementary Lemma 16 (F1* + F2*).

*Using the same assumptions as in Lemma 7, there are no breaks of length* > *m*^1/2^ *with probability greater than* 1 – 3/*n in an optimal chain for large enough n*.

*Proof.* In Section D.3, the structure of the problem does not change. The only difference is the bounds on gap size and spurious k-mers. Asymptotically the homologous gap size is still the same, and both the number of spurious anchors and homologous anchors are smaller by a factor of 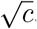, which does not affect the inequalities. Thus all of the bounds in Section D.3 hold except with slightly different probabilities since F1* holds with probability ≥ 1 – 2/*n* instead of 1 – 1/*n*.

#### Supplementary Lemma 17.

*Using the same assumptions as in Lemma 7, the expected recoverability of any optimal chain is* 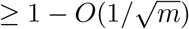 *for large enough n*.

*Proof.* The proof is the same as the proof of Corollary 3 in Section D.4 except we use 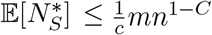 and the failure probability is ≥ 1 – 3/*n*. This doesn’t change the main 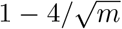 term, so the same big *O* bound holds.

#### Supplementary Lemma 18.

*Under the assumptions of Lemma 7, the expected value of runtime of extension through non-homologous gaps is* 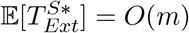.

*Proof.* The proof is the same as the proof of Lemma 9 in Section D.6 except the sketched maximum non-homologous gap length is *m*^1/2^ + *g*‱(*n*). This is still *O*(*m*^1/2^), and 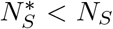, so the expected value of the non-homologous runtime does not change.

#### Theorem 2

*In addition to the hypotheses outlined in Theorem 1, let* 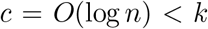 *and* 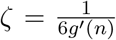 *instead where* 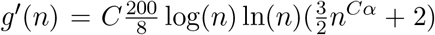. *For open syncmer sketched seed-chain-extend, the expected running time is* 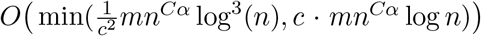 *for extension and* 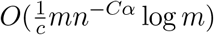 *for chaining. The expected recoverability of any optimal chain is* 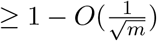.

*Proof.* The expected runtime of chaining and recoverability follows from Theorem 8 and Supplementary Lemma 17. Lemma 13 and Supplementary Lemma 18 give the runtimes under our conditional event space F1*, F2*, which occurs with probability ≥ 1 – 3/*n*, so the same argument as in the proof of Theorem 1 gives the result.

## F Additional figures and tables

**Supplementary Figure 7:**
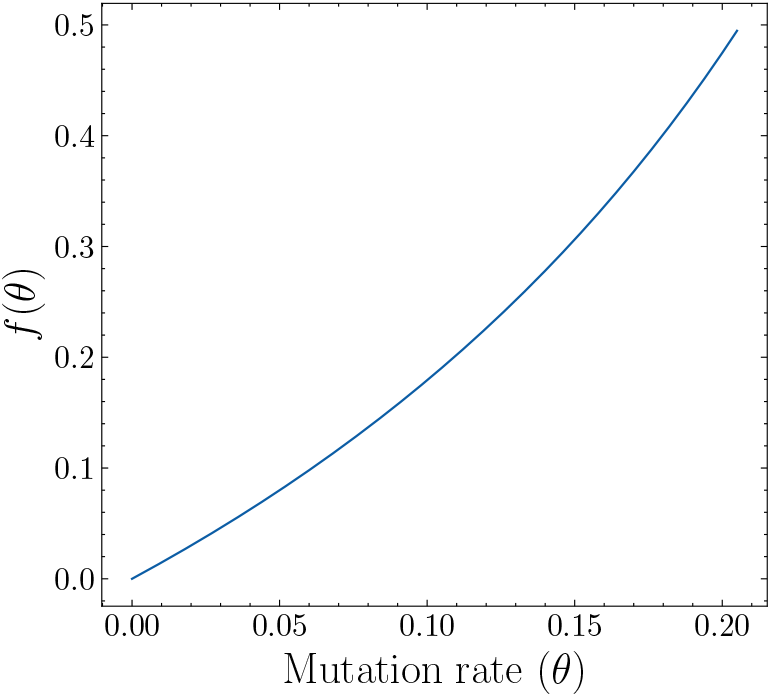
The function *f*(*θ*) for the runtime *O*(*mn*^*f*(*θ*)^ log *n*). Technically, 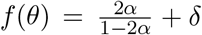 for any *δ* > 0, so we plot 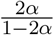 where *α* is a function of *θ* defined as *α* = – log_4_(1 – *θ*). This function is convex on [0, 0.206] so 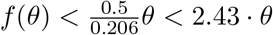.

**Supplementary Figure 8:**
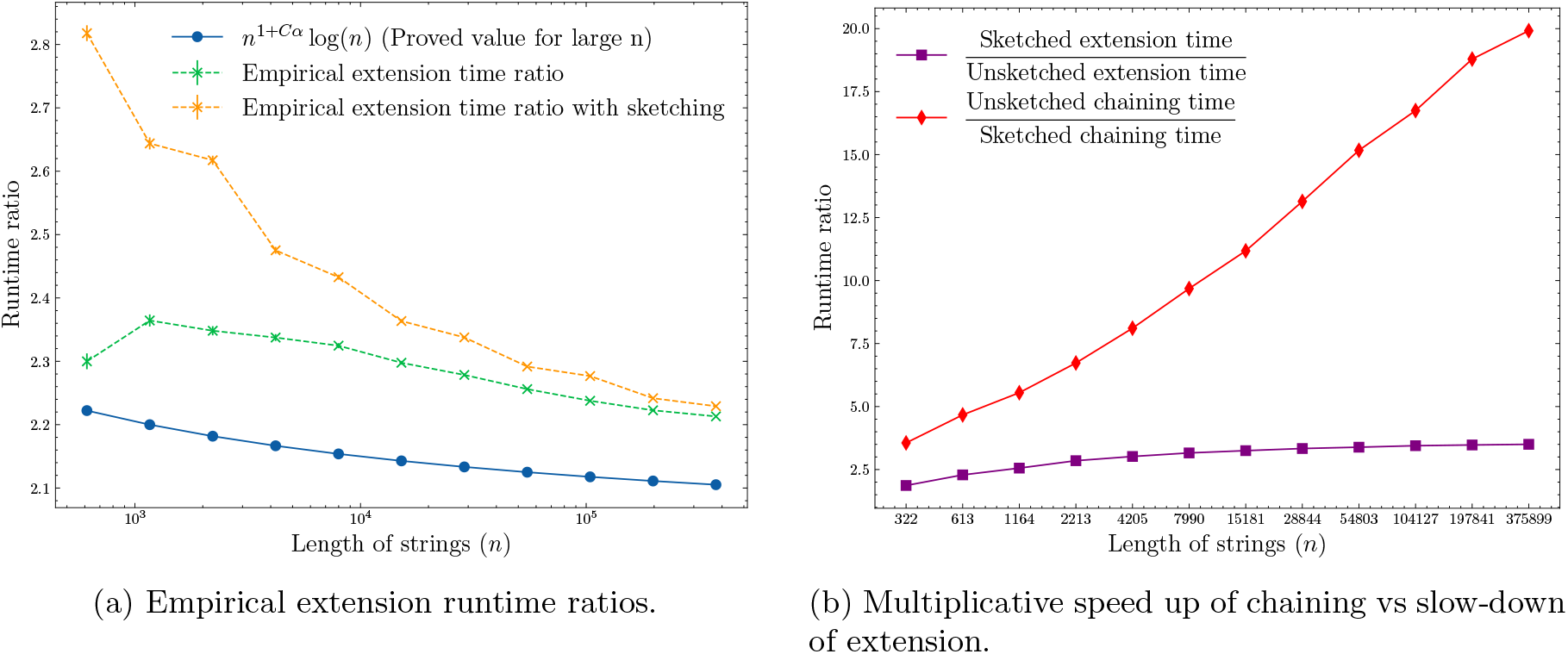
The same experiment as in Figure 1 (a) but with *θ* = 0.05.

**Supplementary Figure 9:**
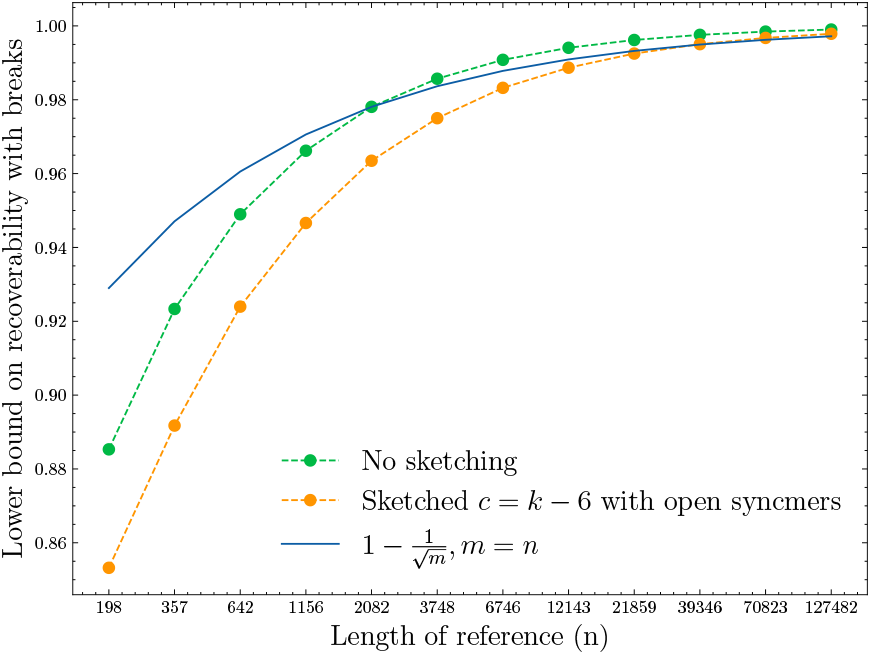
The recoverability of our alignments of two length *n* strings over 50,000 iterations as a function of sequence length *n* where *θ* = 0.10 and *k* was chosen as described in section “Simulated genome alignment experiments”. Breaks were uncommon and most of the recoverability loss is from the (*j_u_* – *j*_1_) term in the recoverability definition, i.e. the chain length being smaller than the sequence length. The lower bound on recoverability is 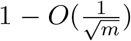 where in Theorem 1 (*m* = *n* in this case), and we plotted 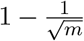 as a non-asymptotic proxy for the asymptotic bound.

**Supplementary Figure 10:**
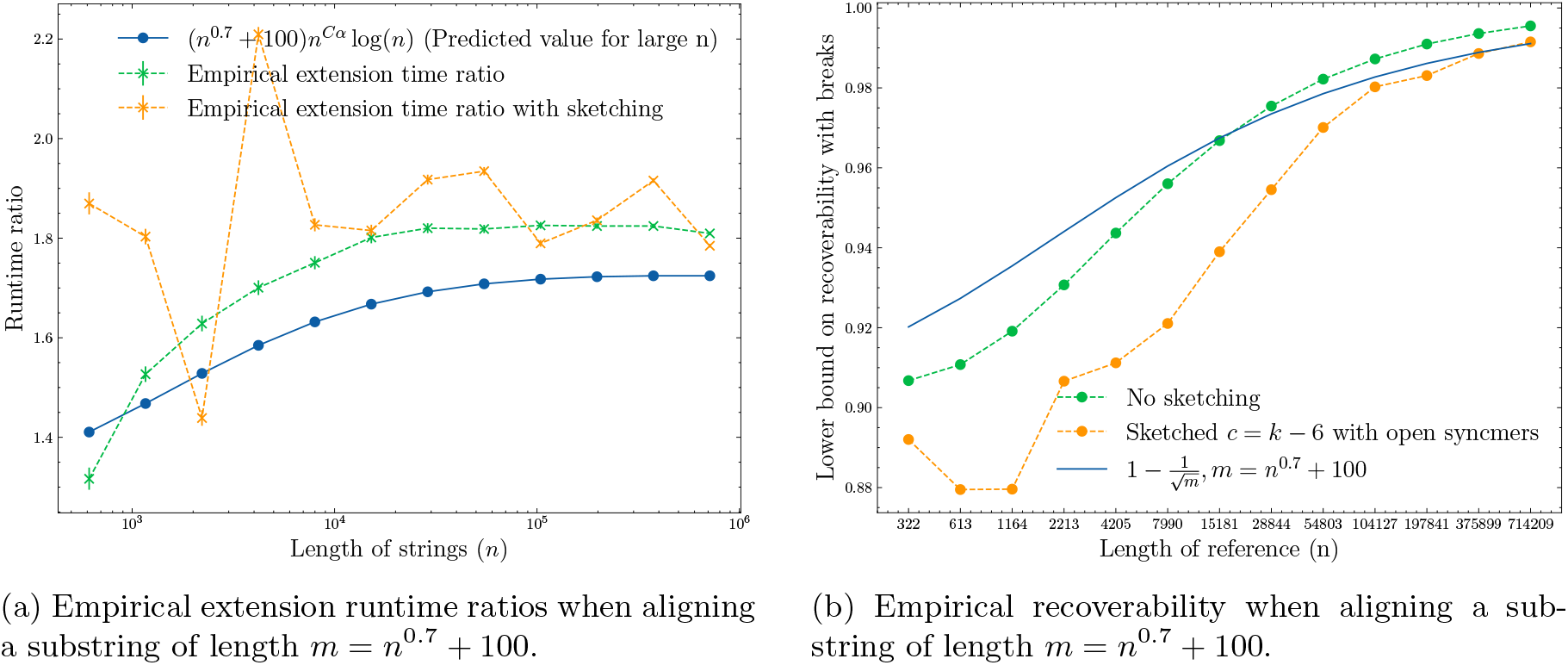
Empirical runtimes and recoverability with *θ* = 0.05 but aligning a mutated substring of length *m* where *m* < *n*. We let *m* = *n*^0.7^ + 100 vary as a function of *n*.

**Supplementary Table 1:** Data sets used for the real nanopore data alignment experiments. Reads were randomly subsampled to the specified number when data sets were too large. We subsampled to 500k reads for the human data set because the reads were more erroneous and had a wider length distribution.

| Genome Name | Genome ID | SRA Read Accession/URL<br>link | Subsampling |
| --- | --- | --- | --- |
| SARS-CoV-2 | NC_045512.2 | SRR22765637 | None |
| Escherichia coli | NC_007779.1 | DRR198814 | 100k |
| Magnaporthe oryzae | CM001231.1 | ERR9878936 | 100k |
| Drosophila melanogaster | NC_004354.4 | SRR21160205 | 100k |
| Homo sapiens | GCA_009914755.3 | https://s3-us-west-2.<br>amazonaws.com/<br>human-pangenomics/index.<br>html?prefix=T2T/CHM13/<br>nanopore/rel3/ | 500k |

## Notes

https://github.com/bluenote-1577/basic_seed_chainer

https://github.com/bluenote-1577/sce-aligner/

### Competing Interest Statement

The authors have declared no competing interest.

### Summary of Updates

Added real experiments. Fixed some typos in the proofs as well. Changed formatting.

